# Conformational dynamics and multi-modal interaction of Paxillin with the Focal Adhesion Targeting Domain

**DOI:** 10.1101/2025.01.01.630265

**Authors:** Supriyo Bhattacharya, Yanan He, Yihong Chen, Atish Mohanty, Alexander Grishaev, Prakash Kulkarni, Ravi Salgia, John Orban

**Author notes:** These authors contributed equally (Supriyo Bhattacharya, Yanan He, Yihong Chen).

## Abstract

Paxillin (PXN) and focal adhesion kinase (FAK) are two major components of the focal adhesion complex, a multiprotein structure linking the intracellular cytoskeleton to the cell exterior. The interaction between the disordered N-terminal domain of PXN and the C-terminal targeting domain of FAK (FAT) is necessary and sufficient for localizing FAK to focal adhesions. Furthermore, PXN serves as a platform for recruiting other proteins that together control the dynamic changes needed for cell migration and survival. Here, we show that the PXN N-domain undergoes significant compaction upon FAT binding, forming a 48-kDa multi-modal complex with four major interconverting states. Although the complex is flexible, each state has unique sets of contacts involving disordered regions that are both highly represented in ensembles and conserved. PXN being a hub protein, the results provide a structural basis for understanding how shifts in the multi-state equilibrium (e.g. through ligand binding and phosphorylation) may rewire cellular networks leading to phenotypic changes.

**Teaser:** The fuzzy complex between Paxillin and Focal adhesion targeting domain is resolved, with functional and translational implications.

## Introduction

Focal adhesions (FAs) are multiprotein structures that link the intracellular cytoskeleton to the cell exterior. These adhesive complexes (FA complexes) are large macromolecular ensembles through which mechanical force and regulatory signals are transduced between the cell and its extracellular matrix(*1*). A critical function of the FA complexes is to facilitate precise spatiotemporal regulation and integration of multiple signaling pathways to output an optimal cellular response to changes in the environment. Thus, the formation and remodeling of focal contacts is a highly dynamic process(*2*) and is modulated by kinases and small GTPases of the Rho family(*3–5*).

Focal adhesion kinase (FAK) and Paxillin (PXN) interact directly with one another(*6*), and constitute the two main components of the FA complex(*3, 7–9*). FAK is a 125 kDa non-receptor kinase that is evolutionarily conserved(*10*), consisting of a central catalytic domain flanked by a large N-terminal domain and a C-terminal region rich in protein–protein interaction sites immediately adjacent to the focal adhesion-targeting (FAT) domain(*3, 11*). The FAT domain is a 125 amino acid four-helix bundle that interacts with PXN, and is necessary and sufficient for localizing FAK to focal adhesions(*2, 10, 12–15*). Through its interactions primarily with FAT, PXN acts as a hub protein and serves as a platform for the recruitment of numerous regulatory and structural proteins that together control the dynamic changes in cell adhesion, cytoskeletal reorganization, and gene expression necessary for cell migration and survival(*16–18*). Furthermore, both phosphorylation and mutation of PXN regulate drug resistance in human lung cancer cells(*19, 20*).

PXN is highly conserved evolutionarily(*21*) and consistent with its intrinsically disordered ensemble, contains several protein-binding modules that allow it to bind to various structural and signaling molecules(*6, 22, 23*). There are four known splice variants of PXN, α, β, γ, and δ, with the α-isoform being the most widely expressed(*18, 23–25*). The α-isoform is a 557-residue protein that can be divided into two domains (**Fig. 1A**). The C-terminal domain consists of four LIM (double zinc finger) motifs(*13*). LIM domains are involved in protein-protein interactions and function as an anchor to localize the molecule to the plasma membrane(*18*). The PXN N-domain is a 311-residue polypeptide chain containing five leucine/aspartate-rich (LD) motifs termed LD1 through LD5 of approximately 12 residues each that are predicted to be helical and display affinity for a range of proteins(*13, 26–28*). Overall, however, the N-domain is thought to be mostly disordered, with four long linker regions (∼20-126 amino acids) connecting the LD motifs(*29*).

**Fig. 1.**
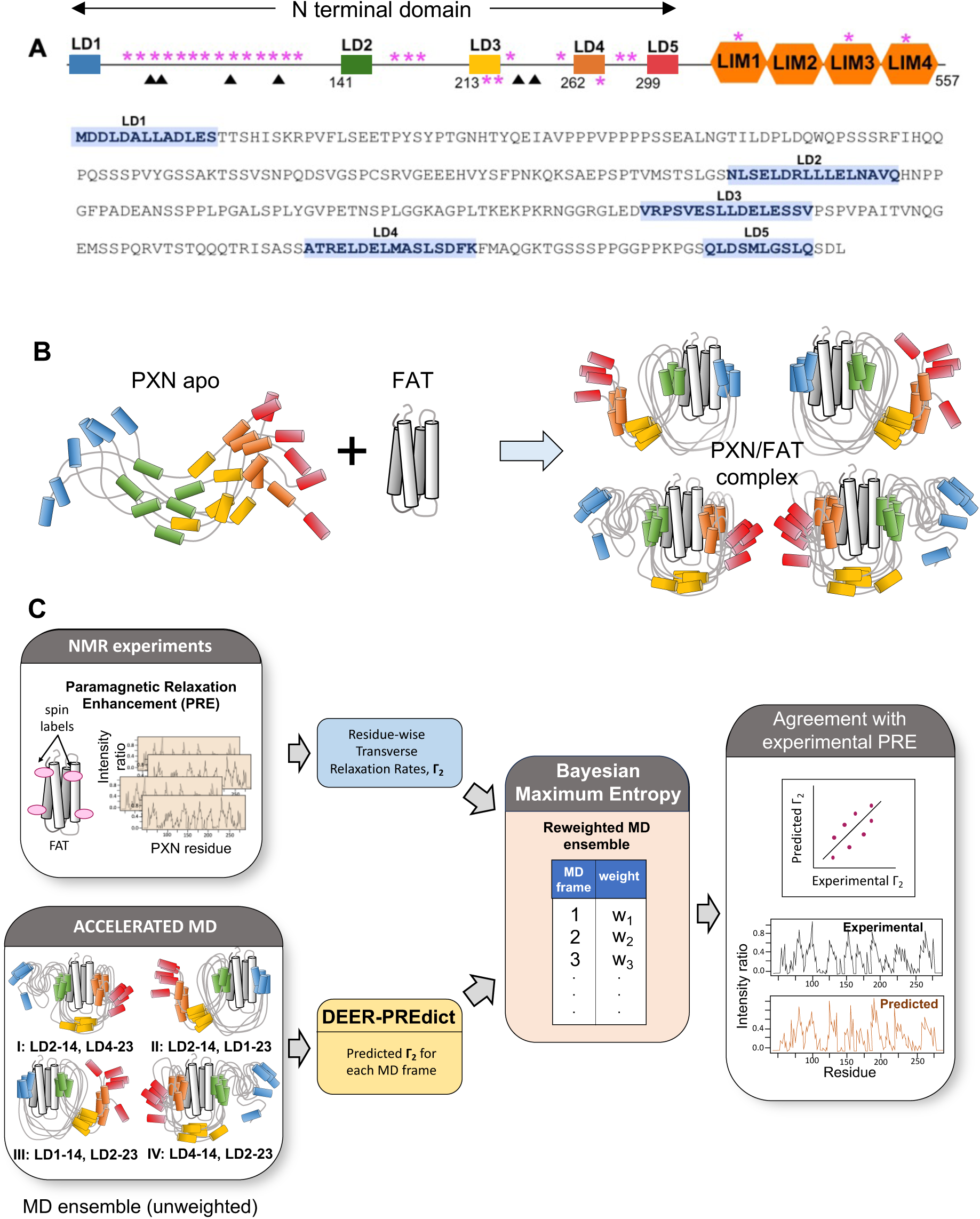
Deriving the conformational ensemble of human PXN bound to FAT. **(A)** Domain organization for the α-isoform of Paxillin. Known phosphorylation (magenta asterisks) and cancer mutation (filled triangles) sites are highlighted. The amino acid sequence for the predicted intrinsically disordered 311-residue N-domain, LD1-5, is shown. **(B)** Schematic describing the multi-modal interaction of PXN with FAT. Disordered regions in the ensemble of PXN conformations (apo and complex states) are represented by grey curves. LD motifs are depicted as cylinders and colored according to the color scheme in panel A. **(C)** In-silico pipeline for deriving multimodal PXN/FAT ensemble starting with MD simulations of the four PXN/FAT bound states, followed by refinement using experimental PRE data.

Previous studies indicated that the binding of PXN to FAT is achieved by the interaction of LD2 and LD4 with two hydrophobic patches (the α1/α4 and α2/α3 sites) on opposite faces of the four-helix bundle of FAT, and that the LD motifs adopt an α-helical conformation upon binding(*2, 10, 12, 14, 29*). Point mutations engineered to specifically disrupt PXN binding to each docking site on the FAT domain individually or in combination revealed that the two PXN-binding sites are not redundant and that both sites are required for FAT function(*10, 30*). X-ray crystallographic structural studies provided insight into how short 10-12 residue peptides corresponding to the LD2 and LD4 motifs of PXN interact with FAT(*2, 10, 12, 14, 30*). However, little is known about the mechanism by which LD motifs engage the FAT domain when present in the context of the entire PXN N-domain.

Here, we employ a combination of NMR spectroscopy, small angle X-ray scattering (SAXS), and molecular dynamics (MD) simulations to characterize the conformational ensemble of the PXN N-domain in its FAT-bound states (**Fig. 1B-C**). Our results demonstrate that, while the N-domain is flexible, it is also conformationally restricted to a significant degree, and that FAT binding leads to further substantial compaction of the PXN N-domain. Our data establish that PXN binding to FAT occurs primarily through three of the five conserved LD motifs, LD1, LD2, and LD4, leading to a dynamic equilibrium between multiple states (**Fig. 1B**). The conformational ensemble and contribution of each PXN/FAT state to the overall equilibrium in this fuzzy complex is described, noting that there are state-specific sets of interactions involving the linker regions between LD motifs. The results thus provide the first detailed characterization of the conformational dynamics for this central interaction in the formation of FA complexes. They also offer a new framework for understanding how post-translational modifications, mutations, and binding with other protein ligands may alter the equilibrium between these differently bound states and impact cellular phenotypic plasticity.

## Results

### NMR assignment of human PXN and FAT

The α-isoform of full-length human PXN, its 311 amino acid N-domain, and human FAT were prepared as described in the Methods section. The two-dimensional ^1^H-^15^N HSQC spectrum of the N-domain has narrow ^1^H shift dispersion, consistent with being intrinsically disordered as predicted (**Fig. 2A**). Moreover, the spectrum is very similar to that of full-length PXN, indicating that the disordered N-domain and the C-terminal LIM domains do not interact significantly in solution (**Fig. S1**). Backbone NMR resonance assignments for the N-domain were obtained using standard triple resonance methods. To help resolve ambiguities and validate assignments, several PXN fragments were also employed, including LD1-2 (residues 1-161), LD2-4 (residues 132-297), and LD2-LD5 (residues 132-311). Most assignments were transferable from the fragments to the N-domain, with minor exceptions for end effects (**Fig. 2A**, **Fig. S2)**. The percentages of main chain amide resonances assigned in LD1-2, LD2-4, and the N-domain are 89.9% (125/139), 90.9% (130/143), and 87.4% (236/270), respectively. Backbone H_N_, N, Cα, Cβ, and CO assignments were deposited in the Biological Magnetic Resonance Data Bank (BMRB) with the following accession codes for LD1-2 (51553), LD2-4 (51554), and N-domain (51555). Backbone resonances were also assigned for human FAT (51556). A chemical shift based CSRosetta structure exhibited a 4-helical bundle fold that closely matched the X-ray structure of human FAT(*31*) (**Fig. S3, Table S1**).

**Fig. 2.**
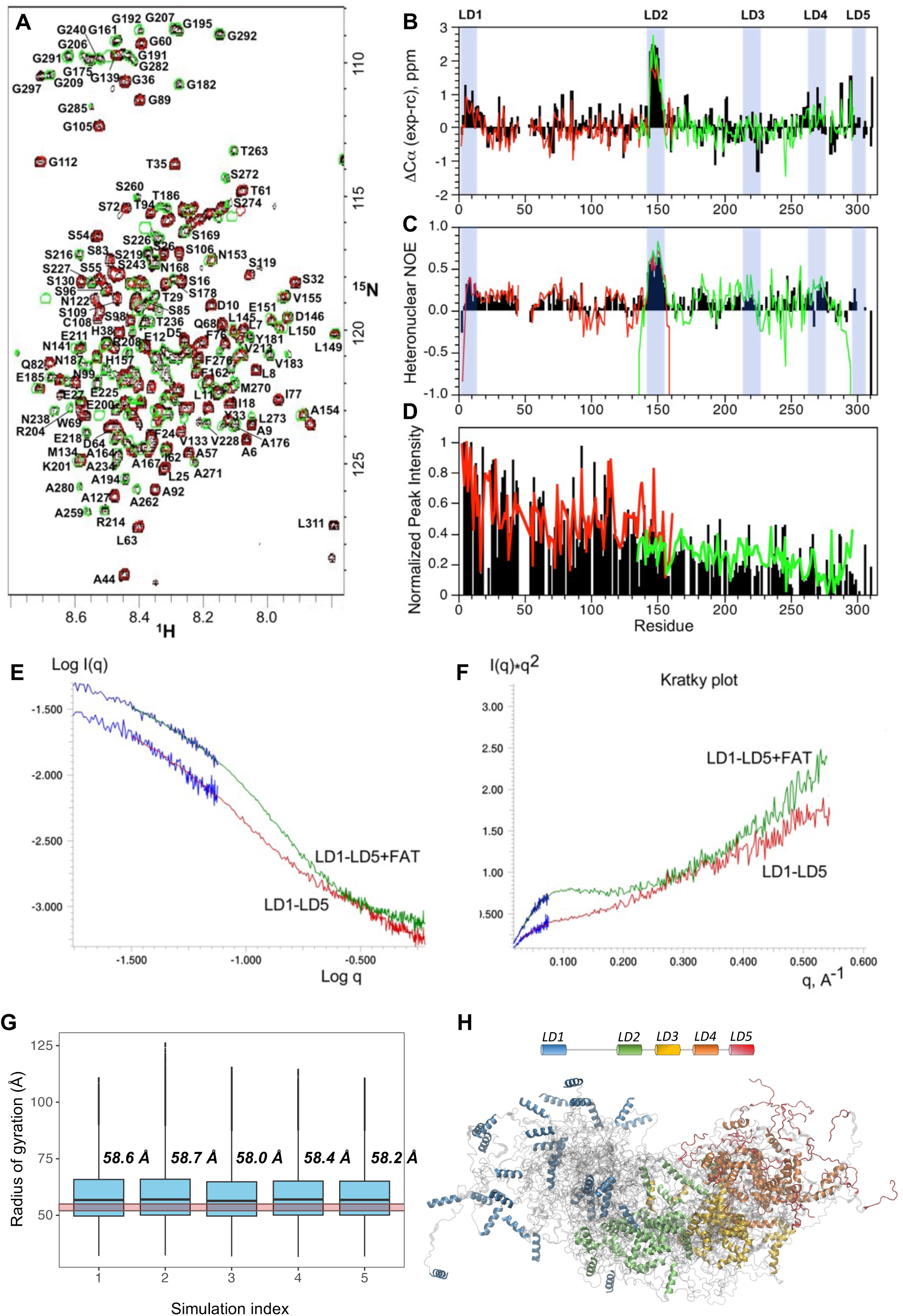
Conformational dynamics of the human Paxillin N-domain and it’s compaction upon binding FAT. **(A)** Two dimensional ^1^H-^15^N HSQC spectrum of the PXN LD1-5 N-domain (black) with main chain backbone amide assignments. Overlaid spectra of the LD1-2 (red) and LD2-4 (green) fragments are also displayed. The largest differences between the shorter fragments and their corresponding regions in LD1-5 are due to end effects. **(B)** Secondary ΔCα shift analysis. **(C)** Steady-state {^1^H}-^15^N heteronuclear NOE values at 600 MHz. **(D)** Normalized peak intensities. The color scheme for (B-D) is N-domain (black), LD1-2 (red), and LD2-4 (green). (**E**) Experimental X-ray scattering data for the PXN N-domain (red), its 1:1 complex with FAT (green), and lowest-angle scattering data acquired with the longest sample/detector distances (blue). (**F**) Kratky plots indicate higher degree of conformational disorder for PXN N-domain compared to its complex with FAT, as evidenced by a pronounced maximum in the complex data at q∼0.1A^-1^, typically associated with folded protein conformations. **(G)** Radii of gyration observed in the AWSEM MD derived PXN conformational ensembles from five independent simulations. Boxes represent the interquartile ranges, while the whiskers represent the two extreme quartiles. Outliers are marked by dots. The experimental radius of gyration (52-55 Å) determined from SAXS is highlighted by the translucent red horizontal band. The average *R*_g_ for each simulation is given above the corresponding box. **(H)** PXN conformational ensemble as obtained from the MD simulations. The LD regions are colored with the color code given in the schematic at the top.

### Conformational analysis of the PXN N-domain

While mostly disordered, the PXN N-domain contains a number of regions within the chain that have distinct secondary structure preferences based on chemical shift analysis. Of these, the LD2 motif has significant helical content (∼80%) based on ΔCα chemical shifts, followed by LD1 (∼30%), and LD4 (∼20%). In contrast, the LD3 and LD5 motifs do not exhibit any appreciable helical character in their unbound states in the context of the entire N-domain (**Fig. 2B**). Consistent with the chemical shift data, the LD2 motif is the least dynamic on the ps-ns timescale with average {^1^H}-^15^N steady state heteronuclear NOE (hetNOE) values of ∼0.7 that are only slightly lower than those for well-ordered helices in globular protein structures (**Fig. 2C**). By comparison, the LD1 and LD4 motifs have lower hetNOEs of 0.3-0.4, in agreement with their diminished helical content and more flexible backbones. Other regions in between LD motifs generally have hetNOE values that range between 0.3 to -0.5, indicating highly dynamic backbone motions on the ps-ns timescale. Additionally, main chain amide resonances for residues ∼150-311 have decreased peak intensities relative to residues ∼1-150 of the N-domain, suggesting exchange between multiple conformational states on a slower timescale (∼μs-ms) for the C-terminal half of the polypeptide chain (**Fig. 2D**). Furthermore, SAXS measurements indicate a radius of gyration (*R*_g_) of approximately 52-55 Å for the N-domain (**Fig. 2E, F**), which is significantly smaller than would be expected for a random coil chain of 311 amino acids (*R*_g_ ∼100-170 Å) and points to appreciable conformational restriction. To gain more insight into the conformational preferences of the N-domain, we performed unbiased coarse-grain MD simulations using the AWSEM method (see Methods). In total, five replicate MD simulations were performed, starting with random initial velocities. All the MD derived ensembles exhibited *R_g_* in the range 58-59 Å (**Fig. 2G, H**) in support of the SAXS measurement. Visual inspection of snapshots from the MD ensemble show that the most significant long range intra-chain contacts occur in the C-terminal half of the N-domain.

### Multi-modal interaction of PXN with FAT

We next determined how the N-domain interacts with its key binding partner, FAT, utilizing a combination of NMR and SAXS measurements, which were inputs for detailed all-atom MD simulations. Titration experiments between ^15^N-labeled PXN and unlabeled-FAT indicated that binding of LD motifs was largely manifested by decreased peak intensity (**Fig. 3A, B**). Thus, peak intensities of ^15^N-labeled N-domain amide resonances were monitored in two-dimensional ^1^H-^15^N HSQC spectra as a function of increasing unlabeled FAT. These data showed that the backbone amide signals of LD1, LD2, and LD4 are the most significantly affected even at sub-stoichiometric amounts of FAT. Effective binding constants (*K*_D_) to FAT of 17±2, 7±2, and 13±2 mM were obtained for LD1, LD2, and LD4, respectively, in the context of the entire PXN N-domain (**Fig. 3C**). We next identified the FAT binding sites accessible to each LD motif. Typically, when there is only one binding site, CSPs and peak intensities would be sufficient to determine a binding epitope in reciprocal experiments where the unlabeled N-domain would be added to ^15^N-FAT. However, the presence of multiple sites meant that such an approach would not provide a clear indication of where each LD motif interacted with the FAT surface. To overcome this ambiguity, intermolecular paramagnetic relaxation enhancement (PRE) experiments were employed, which enabled detailed identification of the mode of FAT-binding for each LD motif. Site-specific Cys mutations were introduced and derivatized with a stable nitroxide spin label (MTSL) at either the N- or C-termini of LD motifs in the natural abundance PXN chain, and PREs were measured to backbone amide protons in ^15^N-FAT. NMR spectra of ^15^N-FAT bound to the PXN N-domain (residues 1-311) were significantly line-broadened and difficult to interpret. However, use of PXN versions encompassing LD1-2 (residues 1-161), LD2-4 (residues 132-297), and LD2-5 (residues 132-311) enabled unambiguous identification of the FAT binding sites in PXN chains with more than one LD motif. The results show that LD1, LD2, and LD4 are all directly involved in interactions with FAT, each binding to a1/a4 and a2/a3 sites on opposite faces of the FAT surface (**Fig. 4A, B, D; Fig. S4**). In contrast, LD3 and LD5 show only minimal PRE effects to FAT, demonstrating that these two LD motifs are not involved in binding to any significant degree (**Fig. 4C, E; Fig. S4**).

**Fig. 3.**
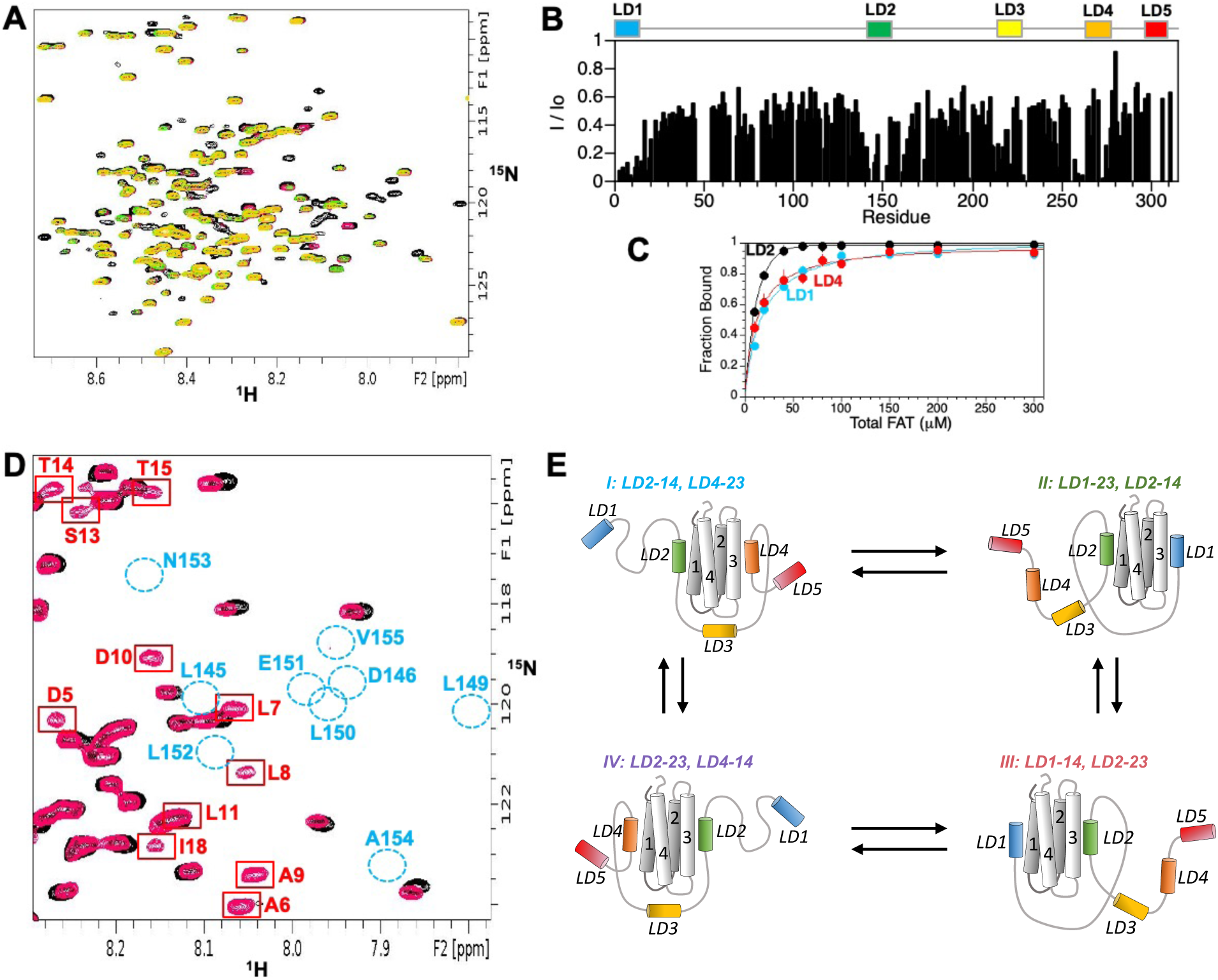
Binding of PXN N-domain to FAT. (**A**) Overlaid two dimensional ^1^H-^15^N HSQC spectra of 100 μM ^15^N-labeled PXN N-domain with increasing amounts of unlabeled FAT added (black, 1:0; red, 1:0.4; green, 1:0.6; orange, 1:1). (**B**) Ratio of FAT-bound amide peak intensity (*I*) to the corresponding intensity in the unbound state (*I_0_*) versus residue for ^15^N-labeled PXN N-domain with 1 molar equivalent of unlabeled FAT added. (**C**) Plots of fraction bound versus the total FAT concentration for LD1, LD2, and LD4 regions in the PXN N-domain. Binding curves were obtained from the decay in amide peak intensities for each region as a function of FAT concentration (see Methods). (**D**) Overlaid two dimensional ^1^H-^15^N HSQC spectra of the 1:1 complex between ^15^N-labeled PXN LD1-2 and unlabeled FAT (black), and the same sample but with 5.45 equivalents unlabeled LD4 peptide added (red). Amide peaks due to the LD1 motif are broadened in the PXN/FAT complex, but gain peak intensity (red boxes) when LD4 peptide is added, consistent with displacement of LD1 from the FAT surface. In contrast, peaks due to LD2 residues (unbound positions shown by blue circles) regain little or no peak intensity upon addition of LD4 peptide. (**E**) Multi-state model of the interaction between the PXN N-domain and FAT under limiting FAT.

**Fig. 4.**
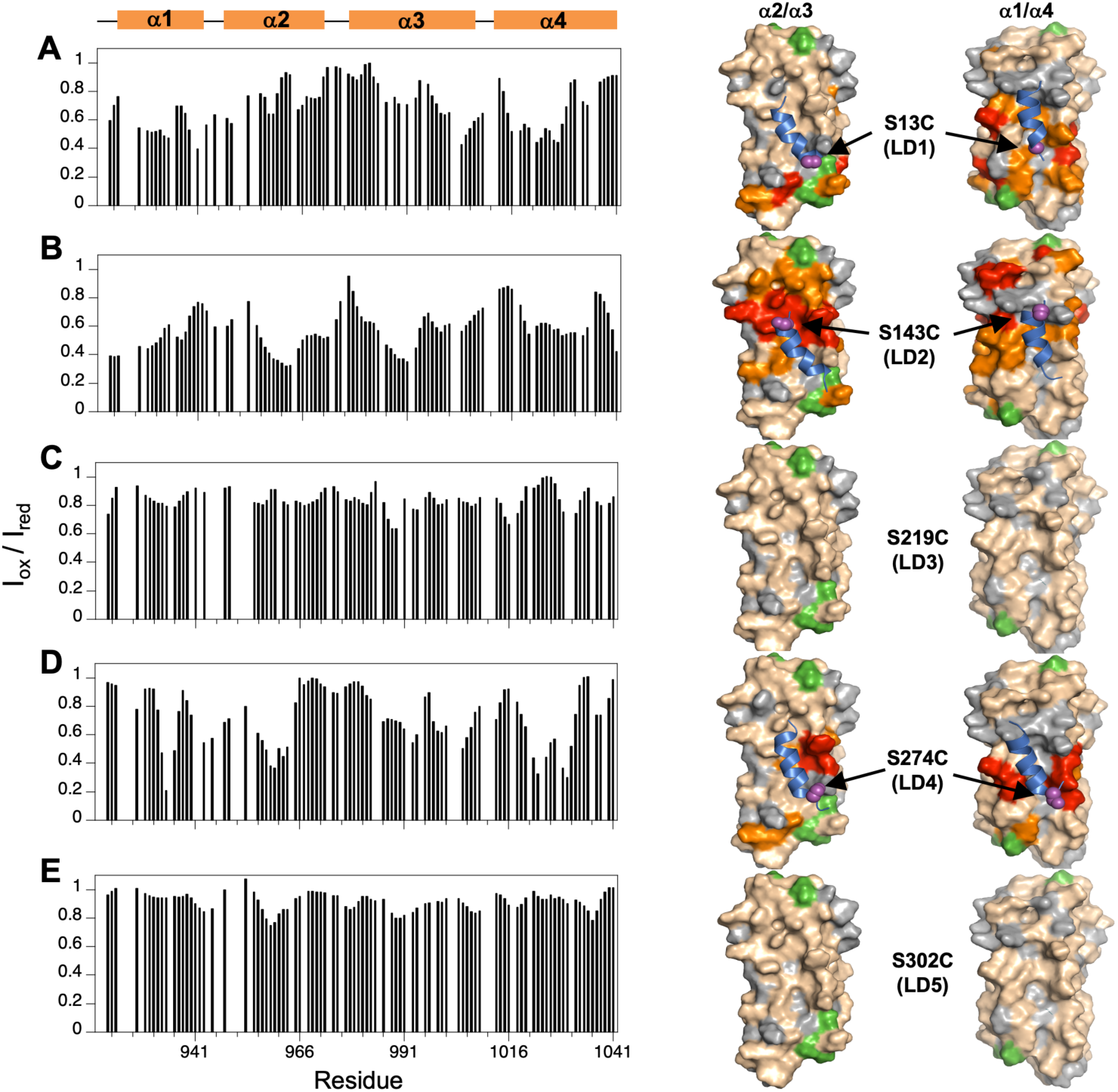
Epitope mapping on the FAT surface using intermolecular PREs indicates that LD1, LD2, and LD4 bind to the α2/α3 and α1/α4 FAT sites. The binding mode of each LD motif to ^15^N-FAT was determined using the following spin-labeled PXNs: (**A**) PXN(LD1-2, S13C-MTSL/C108A); (**B**) PXN(LD2-4, S143C-MTSL); (**C**) PXN(LD2-4, S219C-MTSL); (**D**) PXN(LD2-4, S274C-MTSL); and (**E**) PXN(LD2-5, S302C-MTSL). For each panel, the *I*ox/*I*red plots versus residue are shown (left). Values of *I*ox/*I*red (≤0.5, red; between 0.5-0.6, orange) are mapped onto the surface of FAT showing views for both the α1/α4 and α2/α3 binding sites (right). The PXN chain (blue) is modeled from MD simulations and the attachment site of the spin label in LD1, LD2, and LD4 is indicated. Additional color-coding: Green, proline; Gray, unassigned, overlapped, or exchange broadened signals for which PRE values were not obtained.

In addition to providing binding site information, the PRE data also enabled us to determine the orientation of each LD motif with respect to the FAT surface. Earlier X-ray structures of FAT complexes with short (12-residue) LD peptides(*31*) indicated a preferred orientation of the LD helices such that they are parallel to the α1- and α3-helices of FAT. However, it was not clear if this would also be true in complexes with longer PXN chains encompassing multiple LD motifs. To determine the orientation of LD1 in the context of a longer PXN sequence, the spin label was positioned at the C-terminus of the LD1 motif (S13C-MTSL) in PXN(LD1-2). The clustering of most large PRE effects near one end of the α1/α4 and α2/α3 LD-binding sites on FAT (**Fig. 4A**) indicated that the LD1 helix has a preferred orientation in both sites. Thus, the favored docking configurations have the LD1 helix parallel to the FAT α1- and α3-helices, consistent with the X-ray structures of short peptide/FAT complexes. Similar observations were made in intermolecular PRE experiments where the spin label was placed at the N-terminus of LD2 (S143C-MTSL) (**Fig. 4B**) or at the C-terminus of LD4 (S274C-MTSL) (**Fig. 4D**) in PXN(LD2-4/5). In both these cases, the majority of large PRE effects are grouped near one end of the LD-binding site, indicating that LD2 and LD4 also bind to FAT in a primarily parallel orientation to the α1- and α3-helices of FAT.

We further noted that LD2 displayed the most significant intermolecular PRE effects with FAT (**Fig. 4B**). Moreover, an LD4 peptide added *in trans* to the PXN LD1-2 complex with FAT competes with LD1 for binding to FAT, but does not displace LD2 appreciably over the concentration range used (**Fig. 3D**), suggesting that the LD2 motif acts as the anchoring interaction. This is also consistent with LD2 having the tightest FAT-binding (**Fig. 3C**) and the highest helicity in the unbound state relative to the other LD motifs (**Fig. 2B**). Based on these observations, we built an initial 4-state model in which the LD2 motif maintains binding in all the states (**Fig. 3E**). We then tested this model by comparing the experimental PREs between full-length ^15^N-labeled PXN N-domain and FAT with the calculated PREs derived from MD simulations. The conformational ensembles determined from this model were further validated utilizing SAXS data for the FAT/N-domain complex, which gave an *R*_g_ of 35 Å (**Fig. 2E, F**), demonstrating significant compaction of the disordered PXN chain from its unbound state.

To determine PREs between the PXN N-domain and FAT, the MTSL spin label was attached to specific locations on natural abundance FAT, and PREs to the C108A mutant of the ^15^N-PXN N-domain were measured in FAT/PXN complexes. Four separate sites were designed for attachment of MTSL to FAT. The FAT mutants E984C and Q1006C probed PXN linker conformation around the α2/α3 site (**Fig. 5A, G**), while mutants K1018C and Q1040C probed the α1/α4 site (**Fig. 5D, J**). A clear difference was observed in the PRE profiles for complexes of ^15^N-labeled N-domain with FAT-E984C-MTSL versus FAT-Q1006C-MTSL indicating that, on average, all the linker regions reside nearer to Q1006C when PXN LD motifs are binding to the α2/α3 site (**Fig. 5C, I**, yellow bars). In contrast, PRE effects showed that the PXN linker conformations are more evenly distributed between the K1018C and Q1040C positions around the α1/α4 binding site (**Fig. 5F, L**, yellow bars). The experimental PRE and SAXS observations were then integrated with MD simulations to further quantify the model of the complex between FAT and the PXN N-domain.

**Fig. 5.**
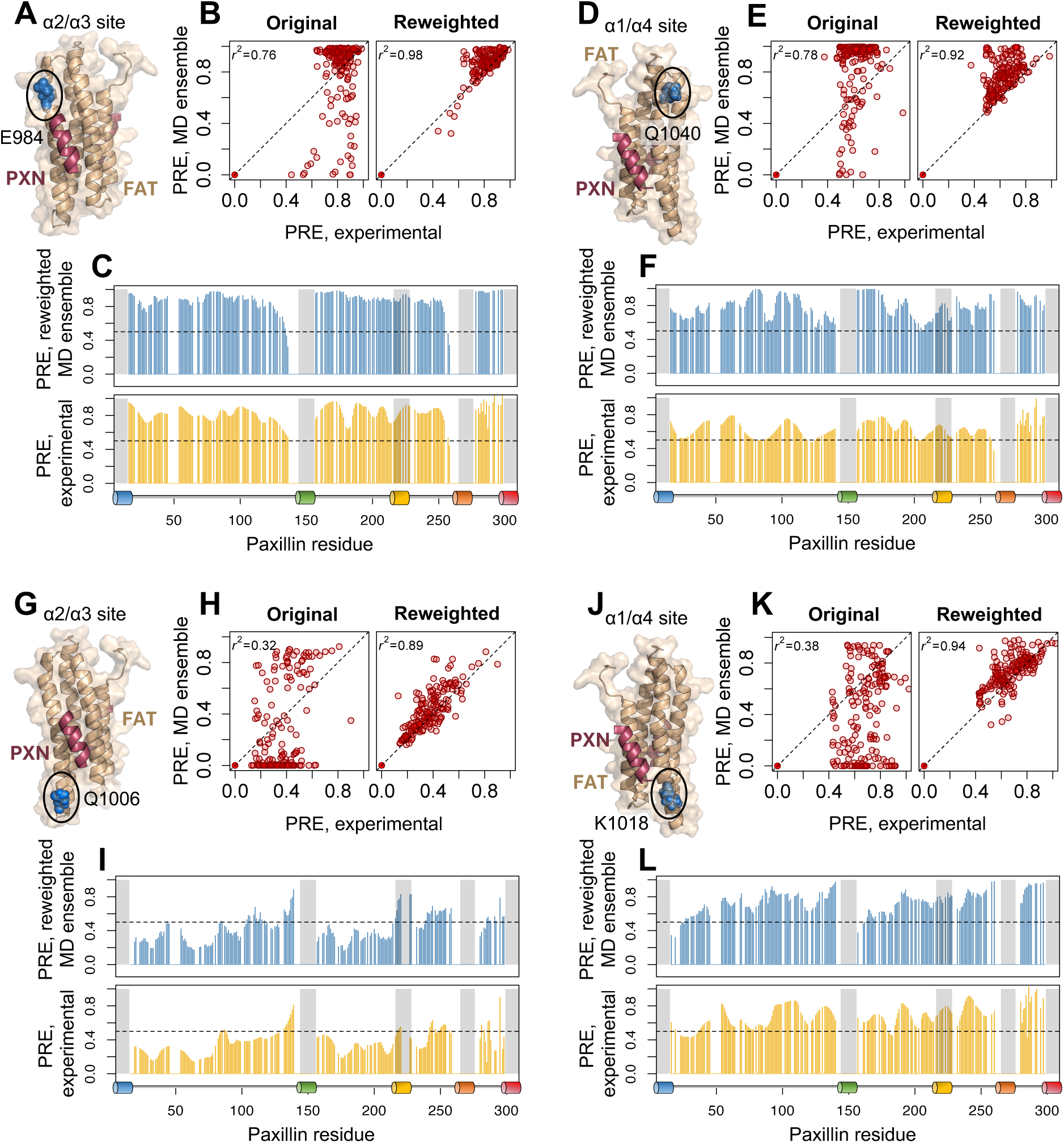
Agreement between experimental and calculated PRE intensity ratios from the MD-derived PXN-FAT ensemble. A stable nitroxide (MTSL) spin label was placed at either **(A)** E984C, **(D)** Q1040C, **(G)** Q1006C, or **(J)** K1018C in the FAT domain (blue, circled) to probe conformational dynamics around either the α2/α3 site or the α1/α4 site of FAT. The position of the LD motif (purple), either LD1, LD2 or LD4, is indicated on the FAT structure. Experimental PREs were measured from the relative peak intensities of PXN backbone amides in the oxidized and reduced states. Reconstructed PREs were calculated from the BME-reweighted MD ensemble, combining the four states to get an ensemble average PRE value for each residue. (**B, E, H, K**) Correlation between experimental and predicted PRE intensity ratios compared among the original and BME reweighted MD ensembles. (**C, F, I, L**) Experimental (yellow) and predicted (blue) PRE intensity ratios are compared along the PXN sequence. Gray bars and colored schematic below the PRE plots indicate the location of LD motifs 1-5. Native numbering for both PXN N-domain and FAT.

### Experimentally guided MD simulations of the PXN N-domain/FAT complex

The PXN/FAT complex was modeled according to the experimental NMR data above, which corresponded with four major configurations (**Fig. 3E**). The initial configuration for each state contributing to the equilibrium distribution of PXN/FAT conformations was constructed using homology modeling. The LD helices and FAT were modelled based on existing crystal structures, and the rest of the PXN linker regions were modelled as flexible loops (details in Methods). Each PXN/FAT configuration was then subjected to coarse grained MD simulations using AWSEM(*32*), followed by all-atom MD simulations using Gaussian accelerated MD (GAMD, see Methods). In total, 5 independent MD trajectories lasting for 1.2-1.4 μs were generated for each PXN/FAT configuration, resulting in 2.8 million conformations. To derive a PXN/FAT ensemble in agreement with the experimental PRE, we reweighted the 2.8 million MD conformations using the Bayesian Maximum Entropy (BME) procedure(*33, 34*) (see Methods). For each MD conformation, the PRE values were calculated using the DEER-PREdict algorithm(*35*). As part of the BME algorithm, the weight for each MD structure (i.e, its contribution towards the equilibrium ensemble) was derived by simultaneously minimizing the error between observed and predicted ensemble-average PRE values, and maximizing an entropy term representing the diversity of the reweighted ensemble. Inclusion of both terms in the cost function achieved improved agreement with experimental PRE, while maintaining a diverse conformational ensemble (*33*). As a result, the structures from MD having higher concordance with experimental PRE were assigned higher weights such that, the average calculated PRE values agreed with the experimental PREs (**Fig. 1C**; Fig. **S7A**). Together, we fitted more than 700 PRE intensity ratios from the four different spin labels to the 2.8 million MD derived structures to obtain an experimentally consistent PXN/FAT ensemble.

To facilitate visualization, the PXN conformations were cast onto the Uniform Manifold Approximation and Projection (UMAP) space using intra-chain distances as similarity measures (**Fig. S9A**) and color-coded according to the weights derived from BME (**Fig. S10A,** see Methods). The four PXN configurations occupied non-overlapping regions in the UMAP, suggesting that they represent distinct intra-chain contacts. Moreover, regions in conformational space having the highest BME weights were localized rather than randomly distributed, indicating that the contribution to the PRE agreement is dominated by specific PXN conformational clusters. All four PXN configurations (**Fig. 6A**) were found to contribute similarly to the PRE agreement, based on their aggregate BME weights (**Fig. 6B**). In agreement with that observation, the clustered regions with high BME weight could be found in all four configurations in the UMAP diagram. The *R*_g_ for each PXN N-domain/FAT configuration calculated using the reweighted ensembles corresponded well with the experimental value from SAXS (**Fig. 6C**).

**Fig. 6.**
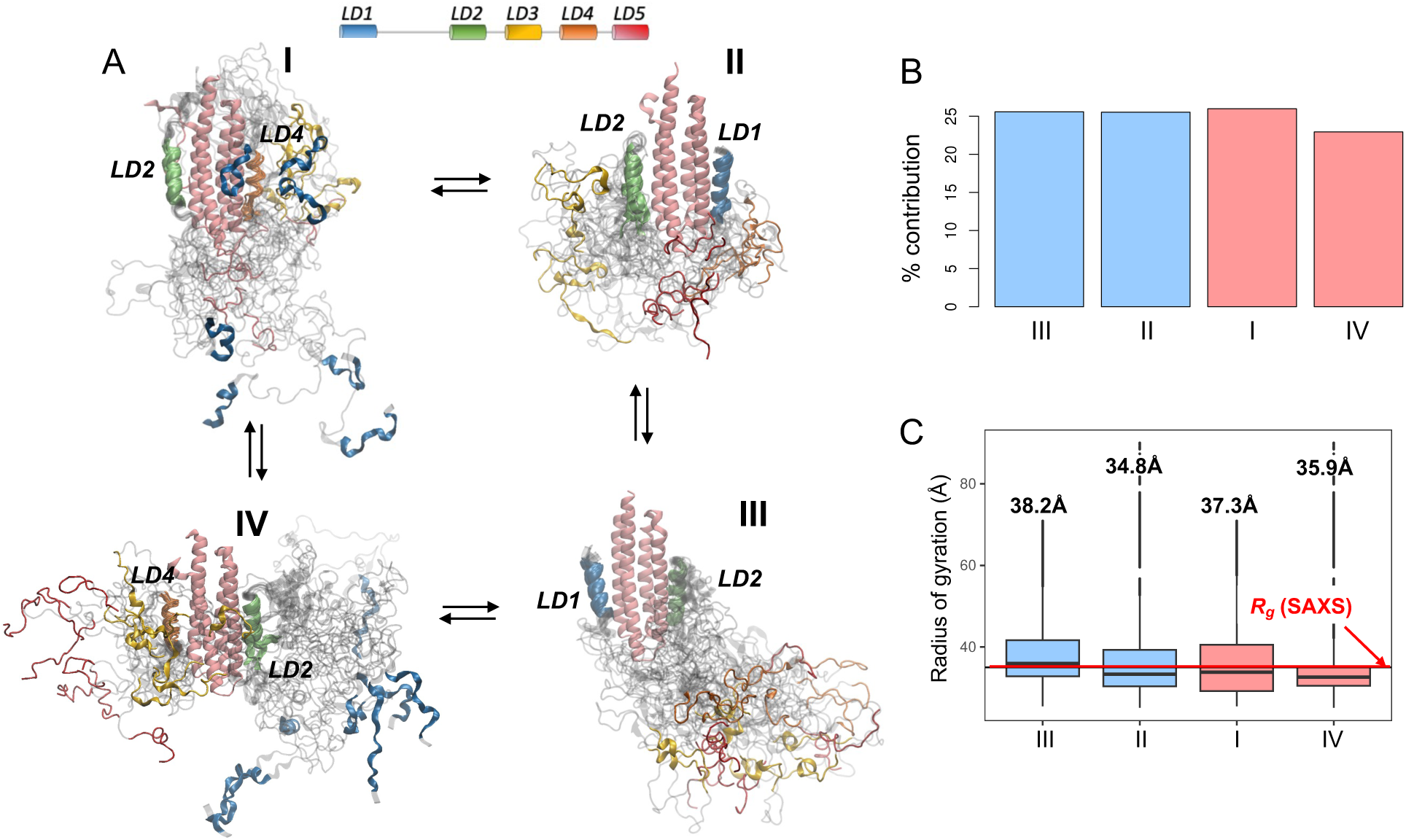
Intermolecular PRE experiments between ^15^N-PXN N-domain and natural abundance FAT enable analysis of the PXN conformational ensemble, including linker regions, relative to the FAT domain. **(A)** Representative structures from the highest contribution cluster for each PXN/FAT state. Individual LD motifs are colored according to the schematic at the top of the panel. The FAT 4-helix bundle is colored in pink. **(B)** Contribution of each PXN state (I-IV) to the experimental PRE intensities, obtained from the BME reweighted MD ensemble of the PXN/FAT complex. **(C)** Box and whisker plot showing the radius of gyration (*R*_g_) calculated from the BME reweighted MD ensemble, for each of the four PXN/FAT states. Boxes represent the interquartile ranges, with the outlier MD conformations plotted along the vertical lines above and below each box. The horizontal red line is the experimental *R*_g_ (35Å) obtained from SAXS.

Comparing the experimental PRE profiles with those derived from MD using the BME reweighted ensembles showed close agreement between the two (**Fig. 5B, E, H, K**, *r^2^*: 0.89-0.98). Moreover, significant improvements in PRE correlation were achieved over the original unweighted ensembles. The re-constructed PRE values re-capitulated many of the general features of the experimental PRE profiles, indicating consistency with our proposed four-state model of the PXN/FAT complex. In particular, the simulated PRE reinforced the experimental observation that there are more significant differences in PRE between the two α2/α3 site probes (**Fig. 5C, I**) compared to the α1/α4 site probes (**Fig. 5F, L**).

We next grouped the FAT-bound PXN MD conformations into 98 clusters (**Fig. S10B**) and the contribution of each cluster to the conformational ensemble was determined (**Fig. 7A**). The highest contribution to experimental PRE agreement came from 23 clusters (**Fig. 7B**), responsible for 87% of the BME-reweighted ensemble. These top clusters are highlighted in the UMAP plot in **Fig. 7C** with representative structures nearest to the cluster centers being shown. One common feature from all these structures is the tendency of the PXN chain to be located towards the “bottom” and “sides” of the FAT, as opposed to the “top”. This agrees with the observation that the PRE probe Q1006C, located near the bottom of the α2/α3 face of FAT (as displayed in **Fig. 5G**), registered the strongest decrease in PRE intensities, indicative of high PXN interaction. Salient features in the experimental PRE profiles can therefore be explained in conjunction with the MD derived structural ensemble, demonstrating the benefit of such an integrative approach.

**Fig. 7.**
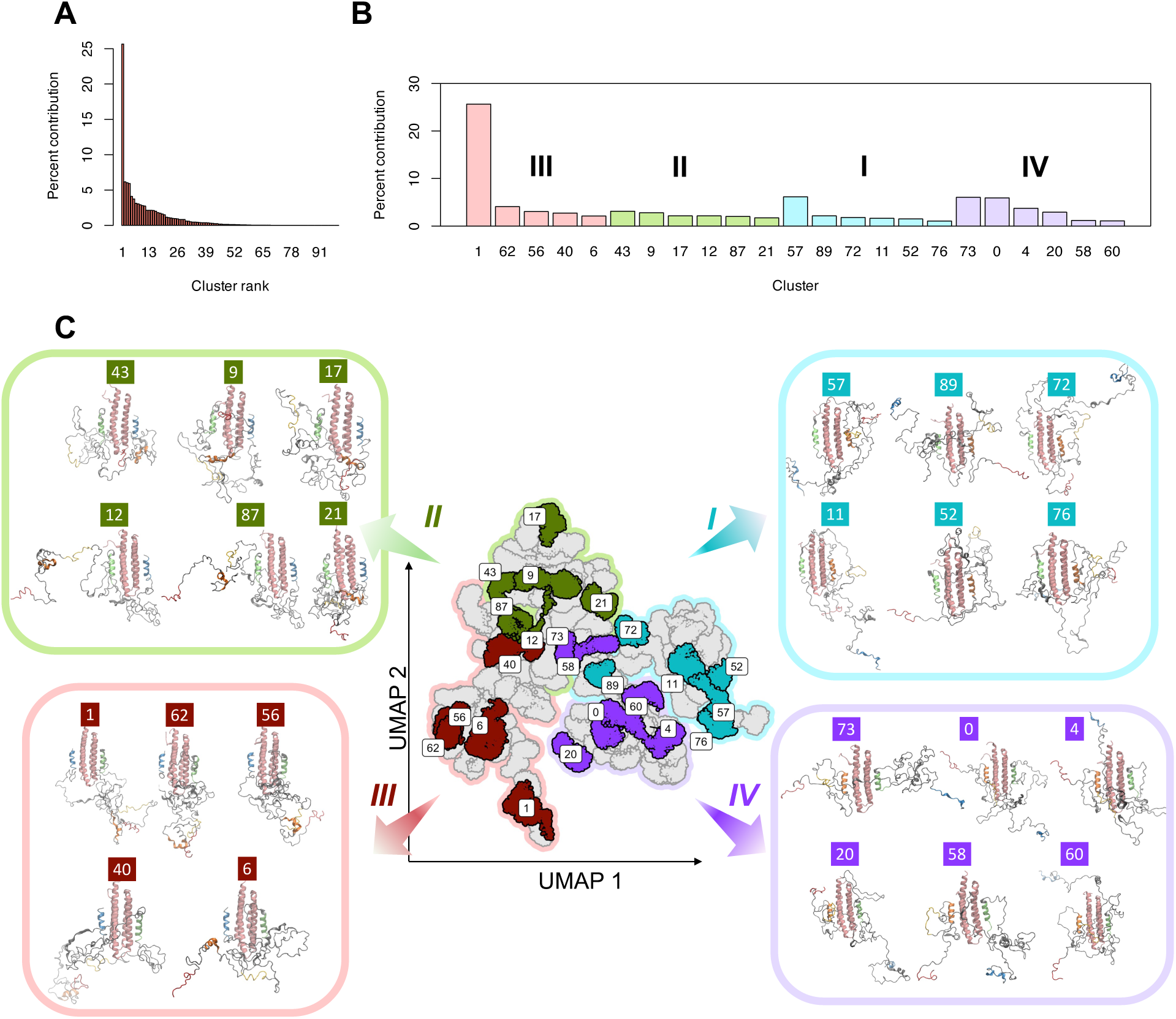
Representation and clustering of PXN N-domain conformations in reduced dimension space using Uniform Manifold Approximation and Projection (UMAP). **(A)** Percentage contributions of all 96 clusters (arranged in decreasing order) towards the conformational ensemble of FAT-bound PXN. **(B)** Percentage contributions of the top clusters (with >1% contribution) from each PXN/FAT state (I-IV) showing highest contribution to experimental agreement with NMR/PRE data. **(C)** Positions of top PXN clusters shown in UMAP space along with the representative structure (cluster centroid using UMAP coordinates) from each cluster. Color coding of the clusters is same as in (B). LD motifs in the representative structures are colored based on the color scheme given in Fig. 1A. FAT is shown as pink helices.

### Each FAT-bound configuration of PXN is characterized by distinct intra-chain contacts

Having established the validity of the MD derived PXN ensembles, we next analyzed the PXN intrachain contacts that are established in the FAT-bound states. Using the top conformational clusters from each PXN/FAT configuration (**Fig. 7C**), we plotted the highest frequency contacts from each of these clusters in a heatmap, organized such that contacts exclusive to each state are clustered together (**Fig. 8A**). This showed distinct PXN intrachain interactions that are highly represented in one PXN/FAT configuration and minimally detected in the other three (red dotted rectangles). Plotting these configuration-specific contacts along the PXN sequence (**Fig. 8B**) showed that many of them were long-range and involved the linker regions between LD motifs, allowing the PXN chain to adopt a compact configuration around FAT.

**Fig. 8.**
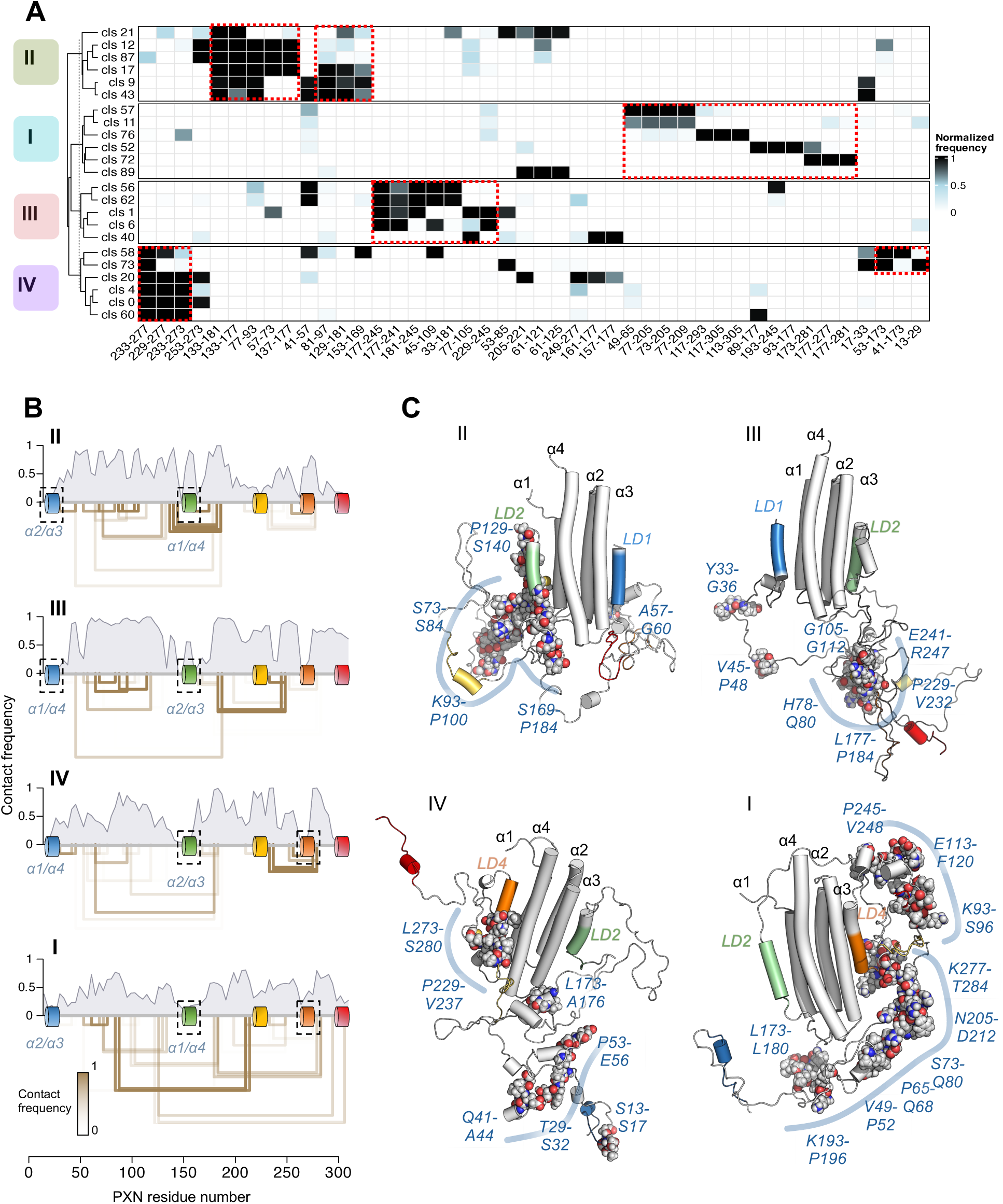
Intra-chain PXN contacts are highly represented and tend to be state-specific. **(A)** Top three most frequently observed inter-segment (see Fig. S9 and S11 for details) contacts among the highest populated clusters (with >1% contribution) from each MD-derived state (I-IV). Cells in the contact map are colored according to the contact frequencies of segment pairs. Short-range contacts between consecutive segments were omitted. Contacts are labeled using the first residue number belonging to each segment. For example, 77-93 indicates the contact between segments spanning residues 77-80 and 93-96. Examples of contacts that are highly represented in one of the four states and minimally detected in the other three are indicated (dashed red boxes). Contacts are arranged such that those exclusive to each state are clustered together in the heatmap. **(B)** Intrachain contact frequency as function of residue number is compared among the four PXN/FAT bound states. The top cluster-specific contacts shown in panel C are represented as open rectangles below each plot, with the termini located near the contacting residues. The height of each rectangle is according to the distance between the contacting residues along the PXN chain, while the color is according to the contact frequency. The locations of the LD helices are highlighted along the x-axis. FAT-contacting LD helices are highlighted in black dashed squares. **(C)** Representative conformations from the four MD-derived states are displayed. For each state, the structure closest to the centroid of the highest populated cluster was chosen as the representative conformation. Helices are shown as cylinders. LD motifs are colored according to the schematic in Fig. 1A. Residue regions corresponding to the highest frequency intra-chain contacts are highlighted as spheres (carbon, gray; oxygen, red; nitrogen, blue).

Analyzing the pairwise intra-chain contact frequency map from the global ensemble incorporating all four configurations (**Fig. S11A**) further confirmed the involvement of linker region interactions. These included contacts between segments comprising residues 81-91 (LD1-LD2 linker) and 281-291 (LD4-LD5 linker), and residues 181-201 (LD2-LD3 linker) and 241-261 (LD3-LD4 linker). Some of these linker region interactions were absent or underrepresented in the contact map from the unweighted ensemble (**Fig. S11B**), even though both contact maps (original and reweighted) shared a common overall topology. Both the original and reweighted ensembles reproduced the experimental radius of gyration, although only the reweighted ensemble showed reasonable agreement with the PRE intensities (**Fig. S12**).

Overlaying the most frequently contacting residues from each PXN/FAT configuration in representative structures (**Fig. 8C**) showed local clusters involving multiple sidechain interactions from residues in the linker regions. Notably, we found tyrosine residues Y31, Y118, and Y181 from known phosphorylation sites to be central partners in these interaction clusters in specific PXN/FAT configurations (i.e. Y181 in II and III; Y118 in I; and Y31 in IV). Together, these results depict a complex multimodal interaction landscape of FAT-bound PXN with potential functional implications (see Discussion).

### PXN-FAT contacts observed in each FAT-bound configuration

Analyzing the contacts between PXN and FAT in the BME reweighted MD ensemble showed persistent interactions involving the LD1, LD2 and LD4 motifs, as expected. However, we also noticed significant interactions with the FAT surface that involved PXN linker regions, particularly LD2-4 and, to a lesser extent, the LD1-2 linker (**Fig. S11C**). The top persistent contacts between PXN linker regions and FAT can be broadly categorized into two groups, involving the α1/α4 and α2/α3 sites of FAT respectively. As seen with PXN intra-chain contacts, each FAT-bound configuration was characterized by distinct state-specific PXN/FAT contacts. While configuration III showed more α2/α3-facing contacts, the other three configurations showed more α1/α4 contacts (**Fig. S11D**). To summarize the involvement of individual PXN and FAT regions in protein-protein interactions, we calculated the interaction probability (fraction of time a given segment is in contact with the other protein) in a sliding window of four amino acid segments along the PXN and FAT sequences for each configuration (**Fig. 9A, B**). Both configurations II and IV showed contacts involving the linker region between LD2-4, with the highest contact frequencies observed in IV, especially involving residues 215-249 (**Fig. 9A**). While some transient interactions were also observed involving the LD1-2 linker (e.g. in configuration I), this region was more heavily engaged in PXN intra-chain interactions, rather than FAT interactions.

**Fig. 9.**
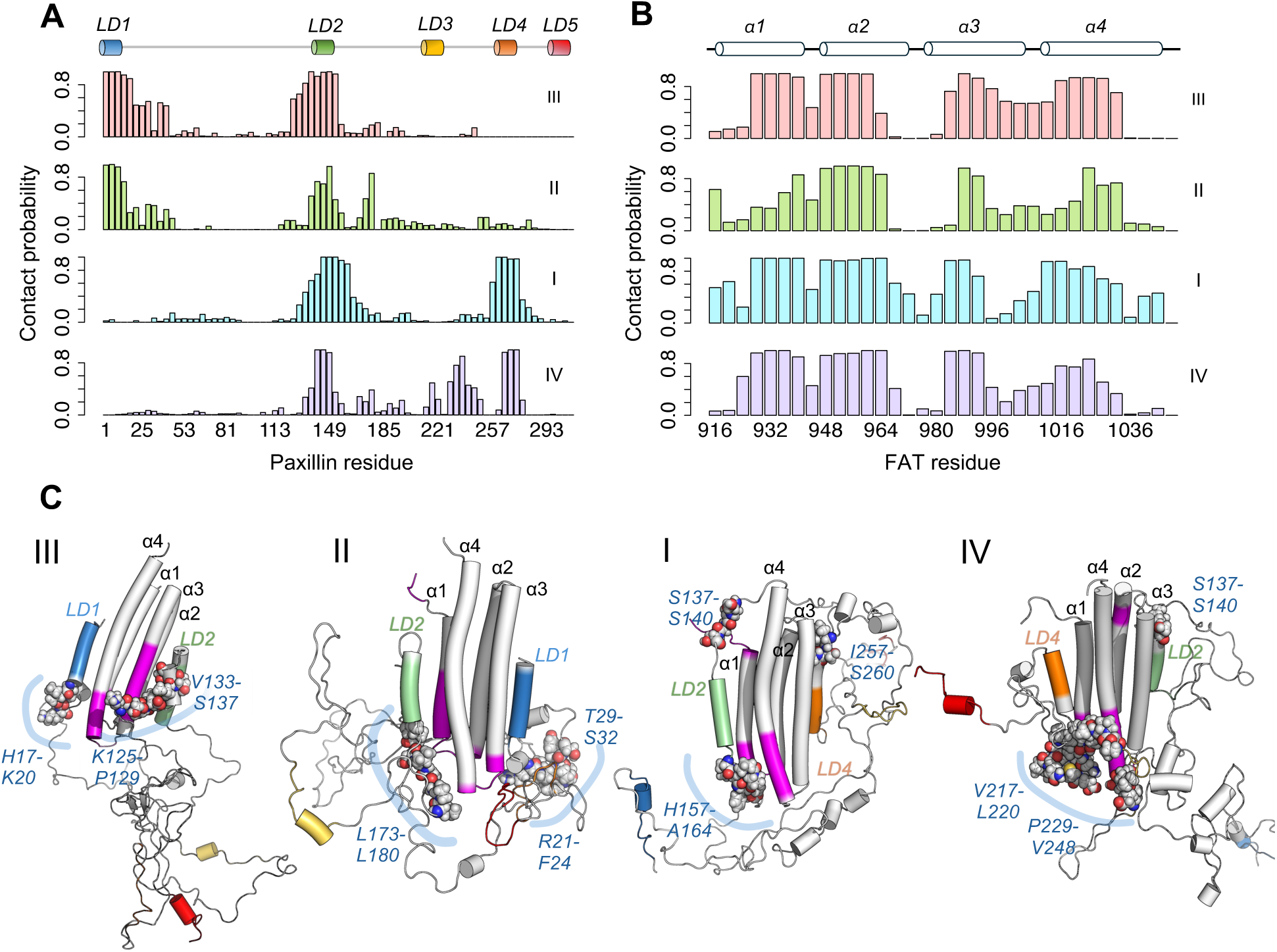
Each PXN state shows distinct FAT contacts. (**A-B**) Intra-chain contact frequencies along the (A) PXN and (B) FAT sequences for the four PXN/FAT states. (**C**) Representative structure from each MD-derived state (see Fig. 9C). PXN residue regions corresponding to the highest FAT contacts unique to each PXN/FAT state are highlighted as spheres. FAT regions corresponding to the highest frequency PXN interactions are colored magenta. LD helices are highlighted according to the color scheme defined in Fig. 1A.

Monitoring the contact frequencies along the FAT sequence indicates distinct differences among the four FAT-bound ensemble configurations (**Fig. 9C**). Overall, the contact frequencies indicate higher involvement of the “lower half” of FAT in contacting PXN, compared to the “upper half”. However, configuration I additionally shows some transient PXN interactions involving the upper half of FAT (**Fig. 9B**) and is supported by the representative structures in **Fig. 9C**.

### Key intra-chain contacts contribute to high conformational entropy of the FAT bound complex

Protein-protein interactions are associated with enthalpy gain due to the formation of intermolecular contacts. This comes at a cost of reduced entropy (entropic penalty) due to conformational restriction upon binding to partner proteins. However, existing evidence indicates that fuzzy complexes involving IDPs maintain their disorder leading to high entropy even in the bound state (*36*), and some of them exhibit high affinity to their binding partners despite being disordered (*37*). This suggests that conformational entropy may play a critical role in stabilizing IDP complexes with their partner proteins (*38, 39*). Yet, the thermodynamic mechanisms (and the sequence regions) contributing to high entropy of fuzzy complexes are not clearly understood.

In this work, we found that PXN undergoes conformational restriction upon binding to FAT, while maintaining significant disorder and mobility in the bound state. To analyze the contributions of the linker regions to the high mobility of the PXN-FAT complex, we applied an information-theoretic approach (**Fig. 10A-B**; Methods) allowing us to estimate the conformational entropy for each PXN residue (**Fig. S140A**)(*40, 41*). Plotting the residue-wise entropy along the PXN sequence revealed several high entropy regions located in the IDR linkers, between LD1 and LD2 and between LD2 and LD4 (**Fig. S14A**).

**Fig. 10.**
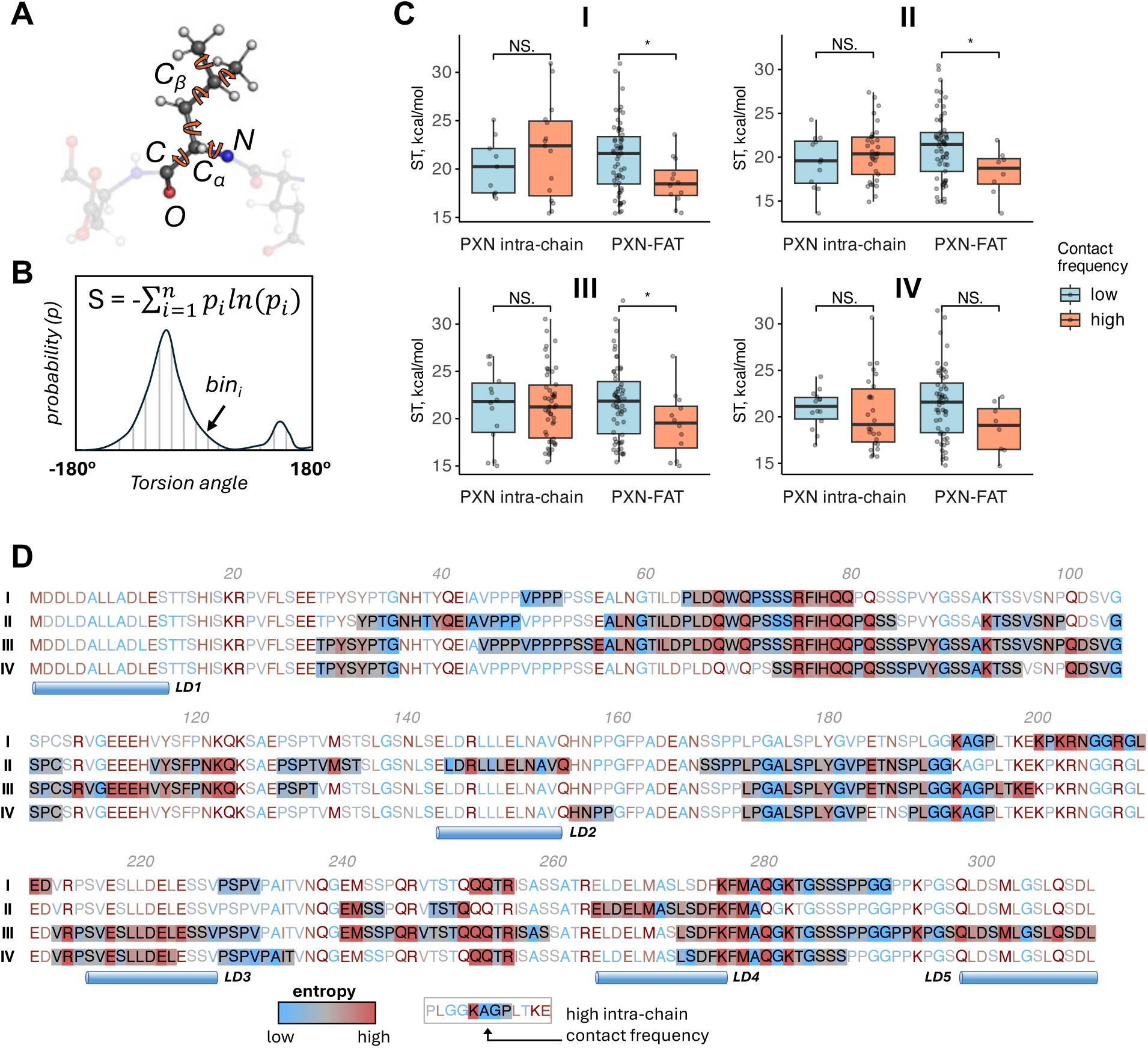
Many PXN intra-chain contacts maintain high entropy while bound to FAT. (**A**) Schematic showing the backbone and sidechain torsion angles in a given amino-acid residue using ball and stick representation; colors – carbon: grey, nitrogen: blue, oxygen: red, hydrogen: white. Individual torsion angles are represented as curved orange arrows. (**B**) Schematic representing a torsion angle probability distribution used in calculating configurational entropy. Probabilities were calculated by dividing the torsion angle range into 35 bins (i.e. bin-width: 10°) and calculating the BME reweighted frequency within each bin. The equation for Shannon entropy is given at the top of the schematic plot. (**C**) Box-plots comparing the entropy of residues showing low (< 20% of time) and high (> 60% of time) intra-chain and PXN-FAT contacts in all four PXN-FAT configurations. Entropy values were converted into energy units by multiplying with RT (R: universal gas constant, T: temperature – 310K). Boxes and whiskers represent the interquartile range (IQR) and 1.5 times IQR respectively. Within each box, individual residues are shown as grey dots. Statistical significance was estimated using Wilcoxon rank-sum test. Pvalues for significance levels were defined as follows - *: 0.01-0.05, NS: > 0.05. (**D**) PXN sequence colored by residue-wise entropy for all four configurations. Regions showing high (>60% of time) intra-chain contact frequency are highlighted with colored backgrounds. Blue cylinders representing the locations of individual LD motifs are shown below.

We further analyzed the entropy of the PXN residues in conjunction with their propensities to form intra-chain and FAT contacts (**Fig. S14A**). In general, contact-forming residues are expected to have low entropy due to their restricted mobility. Peaks in PXN-FAT contact frequency plots (e.g. LD1 in states II and III, LD4 in states I and IV, LD2 in all four states) coincided with low entropy regions in the PXN chain in all four configurations, indicating that PXN residues contributing to FAT binding indeed suffer entropy loss, as might be expected. In contrast, PXN regions with high intra-chain contact frequency did not typically correspond to low entropy regions. In fact, several of the major intra-chain contacts coincided with high peaks in the entropy plots (**Fig. S14A)**. Scatter diagrams of contact frequencies versus entropy further confirmed this observation (**Fig. S14B**). In line with this, high frequency FAT contacts on average were significantly lower in entropy than low frequency contacts (p<0.05) in 3 out of 4 FAT bound configurations, while the entropy differences between high and low intra-chain contacts were insignificant (**Fig. 10C**). Our analysis showed that many PXN intra-chain contacts maintained high entropy in the FAT bound state (e.g. the whiskers of the box plots in **Fig. 10C)**, thereby contributing to the conformational entropy of the PXN-FAT complex. Highlighting the intra-chain contacts along the PXN sequence indicated that the high entropy contacts involved multiple polar residues from the linker regions that were mainly glutamine, along with several lysines, arginines and glutamates (e.g. Q68, Q70, R75-Q80, Q82, K94 between LD1 and LD2; K193, Q253, Q254, R256 between LD2 and LD4; Q281, K283 between LD4 and LD5; **Fig. 10D**). These residues showed consistently high entropy in at least three out of four FAT bound configurations, despite being engaged in high frequency intra-chain interactions.

## Discussion

The large size of the central binding partners in the focal adhesion complex, PXN (551 residues) and FAK (1052 residues), and the highly dynamic nature of their interaction, pose significant challenges for obtaining detailed structural information. These two factors in combination limit the applicability of current structure determination methods to studying the complex in its completely native state. Therefore, we adopted a strategic approach in which the most relevant components to understanding how PXN interacts with FAK were employed. This enabled detailed structural characterization of the complex between the N-domain of PXN (311 residues) and the FAT domain (125 residues), utilizing a combination of solution NMR/PRE and SAXS measurements that are integrated with MD simulations. In particular, PREs provided a powerful method for obtaining information about transient contacts due to their sensitivity over longer distances than NOEs. In addition, the ability to label specific sites with stable nitroxide spin labels simplified analysis of intermolecular contacts significantly, allowing more facile deconvolution of contributions from different flexible states. The results demonstrate that, while the N-domain is largely disordered in its unbound state, it is nevertheless relatively restricted in conformational space and becomes even more so upon interacting with its key binding partner, FAT.

Both components of the PXN/FAT complex have multiple binding sites. The FAT docks LD motifs at α1/α4 and α2/α3 sites, located on opposite faces of its 4-helix bundle fold, while the PXN N-domain binds FAT primarily through its LD1, LD2, and LD4 motifs. This results in a multi-state PXN/FAT complex in which the centrally located LD2 motif plays an anchoring role, consistent with it having the most stable helical structure of all the LD regions in the N-domain. The structural model is therefore of a dynamic equilibrium between four conformational states (I-IV), where the LD2 motif binds at α1/α4 or α2/α3 of FAT while LD1 and LD4 compete for binding to the remaining open site. This model is supported by all-atom MD simulations that reconstruct the experimental PREs and the SAXS-derived *R*_g_ value for the N-domain/FAT complex near quantitatively. While we cannot completely rule out contributions from other states, such as reversed LD orientation in FAT sites, these have low populations and do not contribute significantly to the overall PRE profiles. Although the PXN polypeptide chain undergoes significant compaction upon binding to FAT, each of the four bound states remains relatively flexible, as reflected in their ensemble representations. As such, the interaction between the PXN N-domain and FAT may be considered as an example of a “fuzzy” complex(*42, 43*), with the LD2 contact being the initial hook, and the linker regions exploring transitions between states I-IV utilizing a fly-casting type of mechanism(*44*). Also, we found multiple polar residues in the linker regions maintaining high conformational entropy in the FAT bound state. These residues, despite high intra-chain contact frequency, maintained their entropy and flexibility, presumably through transient interactions with multiple PXN residues. Of note, we found a glutamine-rich region between LD1 and LD2 (residues 68-82) with high entropy intra-chain contacts in multiple FAT bound configurations. These high entropy intra-chain contacting residues are also conserved among the PXN sequences in diverse species, as indicated in **Fig. S13**. In a recent study, glutamine-rich regions in an intrinsically disordered protein were shown to mediate dynamic intra- and intermolecular interactions(*45*). The glutamine-rich PXN regions may therefore have evolved to maintain high entropy in the FAT bound configurations, thereby contributing to the stability of the bound complex.

Despite flexibility in the linker regions between the LD motifs of the FAT-bound PXN chain, there are structural features particular to each bound state that are worth noting. For instance, hydrophobic and aromatic residues in the IDRs form transient but nevertheless state-specific intrachain contacts (**Fig. 8**), which contribute to the compaction of the PXN chain around the FAT domain and the relatively low observed *R*_g_. To further understand the significance of these intramolecular PXN contacts, we looked at PXN sequence conservation among multiple species (**Fig. S13**). This analysis revealed, unsurprisingly, that the LD motifs were near-invariant, but also showed high conservation in some parts of the linker regions between them. Notably, most of the PXN intrachain contacts that are highly represented in each ensemble state (**Fig. 8C**) contained a significant fraction of conserved residues. These intrachain contact regions included, amongst others, H116-F120 in state I, P129-S140 in state II, Y33-G36, V45-P48, P229-V232 and E241-R247 in state III, and Q41-A44, P53-E56 in state IV. Similarly, we noted that each PXN/FAT state had distinguishing intermolecular interactions, and found that the highest probability contacts between PXN linker residues and FAT also tended to correspond with regions of the PXN chain that were well conserved (**Fig. 9, Fig. S13**). These intermolecular contacts included S137-S140 and I257-S260 in state I, R21-S32 and L173-L180 in state II, T29-S32 and K125-P129 in state III, and P229-V248 in state IV. As PXN is a hub protein in the focal adhesion complex, some of the conserved linker regions are likely protein interaction sites. One that is well known is the polyproline site between residues 44-53, which is responsible for interaction with the SH3 domains of many partners such as the muscle fiber protein ponsin(*46*). Moreover, there are approximately 25 known phosphorylation sites in the PXN N-domain(*17, 18*), mostly in the linker regions (**Fig. 1**). We therefore suggest that ligand binding and/or phosphorylation events are likely to affect the equilibrium between PXN/FAT states, which may contribute to specific phenotypes such as cell motility and survival(*47, 48*).

Exploring the conformational space of larger IDPs using MD simulations involves challenges such as limited conformational sampling and force field deficiencies(*49*). In this work, we used the a99sb-disp force field which was designed to reproduce the dynamics of both IDPs and folded proteins. To address the challenge of conformational sampling, we employed GAMD, a special form of accelerated MD, which was shown to significantly improve the sampling of protein conformations in MD simulations (*50*). However, given the size of the PXN chain, it was not feasible to start atomistic MD from an extended conformation, as this would have necessitated a significantly larger simulation box, slowing down the simulations. Initial use of AWSEM coarse-grain MD to collapse the PXN chain around FAT therefore facilitated the subsequent atomistic MD simulations, since the collapsed PXN/FAT structure could be accommodated within a smaller simulation box.

Due to the structural flexibility of IDPs, deriving their conformational ensembles by combining MD simulations with experimental measurements is challenging(*51, 52*). This task becomes more difficult for longer polypeptide chains such as PXN due to the vast degrees of freedom. To address this challenge, we employed BME, which is an established statistical approach for refining MD ensembles using experimental data. This approach is advantageous because it retains the conformational diversity of the derived ensemble, while achieving quantitative agreement with experimental data. Thus, BME is especially suitable for IDPs, where diverse conformations contribute to the ensemble average experimental properties. To facilitate the reweighting process, we used more than 700 PRE intensity ratios from MTSL probes at four different locations on FAT, thereby minimizing the ambiguity of the derived weights. Additionally, technical challenges of fitting PRE intensity ratios to MD ensembles were addressed by providing practical solutions (e.g. by truncating Γ_2_ beyond a cutoff), which will benefit future efforts in deriving ensembles of other IDPs. Notably, the original MD ensemble agreed with the macroscopic dimension of the PXN/FAT complex (experimental *R_g_*), but failed to reproduce the PRE intensities (**Fig. S12A-B**), while the BME reweighted ensemble agreed with both (**Fig. S12C-D**). Refinement based on multiple localized measurements (e.g. PRE, that reflect intermolecular distances) therefore can better resolve the structural nuances within IDP ensembles, especially for larger proteins and complexes.

One limitation of our domain approach is that the potential perturbing role of other regions in PXN and FAK on the multi-state equilibrium is not addressed. For example, the adjacent 120 residue linker region connecting the N-terminal α1-helix of FAT to the FAK kinase domain is not included here. However, our ensemble description of the complex shows that the PXN chain is positioned away from this linker site in all the FAT-bound states, suggesting that the FAT-linker is unlikely to have an appreciable steric effect on transitions between states (**Fig. 7C**; **Fig. 8C**). Our data also indicate that the N- and C-domains of PXN do not interact significantly in the absence of other partners (**Fig. S1**). Nevertheless, possible long-range effects on the complex from other parts of the FAK chain cannot be completely excluded, particularly as its kinase domain can phosphorylate PXN. It should also be noted that the multi-state model is for PXN N-domain binding to stoichiometric levels of FAT. Under these conditions, it has been shown that binding of one LD motif to FAT promotes association of another LD motif from the same PXN molecule with the second FAT site in a cooperative mechanism, preventing interaction of more than one PXN per FAT(*29*). Our experimental data also supports this scenario. Additionally, it is possible that the PXN chain binds to multiple FAT molecules when there is an excess of FAT. Under these conditions, the second FAT would associate with an unbound LD1 or LD4 motif, leading to disruption of the multi-state equilibrium and a potential pathway to hetero-oligomeric states. These types of hetero-oligomeric interactions are not detected here as they would yield significantly more broadened NMR signals and a larger *R*_g_ than seen in the current study. The population of hetero-dimeric versus hetero-oligomeric PXN/FAT states in cells will presumably depend on the relative expression levels of PXN and FAK. Notably, in numerous cancers, PXN is overexpressed relative to FAK, which would suggest a PXN/FAT structural model similar to that described here(*53*).

The multimodal interactions between PXN and FAT may also have significant functional implications. Due to their flexibility, IDPs such as PXN are capable of interacting with multiple partners, signifying their role as hubs of protein interaction networks (PINs) (*54–56*). It has been proposed that modulating the conformational ensembles of IDPs is an evolutionarily conserved mechanism of PIN rewiring, enabling switching of cellular phenotypes in a reversible, dynamic manner (*57, 58*). The current study provides a structural framework for understanding how this may work in FA complexes.

Firstly, the multimodal PXN-FAT interaction exposes specific protein binding motifs in the PXN chain in a state specific manner, facilitating the binding of distinct partner proteins. The heterogeneity of the FAT-bound PXN ensemble could therefore be exploited by the cellular machinery to regulate the FA composition or downstream signaling based on environmental cues (*59, 60*). This finetuning may be achievable via altered phosphorylation of PXN by specific kinases, as discussed above.

Secondly, both PXN and FAK have been shown to sequester key proteins at the FA by undergoing liquid-liquid phase separation (LLPS) via oligomeric complexation (*61*). Previous studies have implicated the role of hydrophobic residues in IDR sequences for LLPS formation (*62, 63*). Notably, in this study, we have found that hydrophobic residues in PXN form local clusters of intra-chain contacts that are state-specific. Modulation of these local interaction hubs through equilibrium shifts among the PXN-FAT states (e.g. via phosphorylation) could have a significant impact on PXN mediated LLPS, with far-reaching consequences on cellular behavior. Taken together, the malleability of the multimodal interaction dynamics of PXN may be harnessed by the cell for flexible decision making in response to extrinsic signals or environmental changes.

In summary, the results presented here provide the most extensive molecular description to date for the highly dynamic interaction between PXN and FAK, two critical components of the focal adhesion complex. They also offer a basis for further studies aimed at determining how perturbations, such as phosphorylation, ligand binding, and mutations, might lead to different structural and functional outcomes. Moreover, the importance of the PXN/FAT interaction in both cancer metastasis and drug resistance mechanisms suggests that it may be a viable therapeutic target despite its flexibility(*64*) as was recently supported (*65, 66*). Indeed, an in-depth understanding of the multimodal interaction between PXN and the FAT domain of FAK may facilitate the design of therapeutics that can disrupt this interaction in novel ways. Small molecules may be able to shift the equilibrium between different PXN/FAT states (for example), particularly since state-specific interactions are a characteristic of the multi-state equilibrium. The results therefore have both fundamental relevance as well as possible translational applications. Given the multiple roles of PXN in development(*67*), cell migration(*53*), and angiogenesis(*53, 68*), our study should pave the way for more detailed investigations on its ensemble-function relationships. Additionally, our approach addresses multiple practical considerations in deriving structural ensembles of large flexible complexes using MD-based methods, which may have general applicability.

## Materials and Methods

### Sample preparation

PXN and FAT genes and gene fragments were cloned into an eXact tag pH720 vector system. This approach places an albumin-binding (GA) domain followed by the prodomain of subtilisin at the N-terminus of the target sequence. The expressed protein was then purified on an engineered subtilisin column, followed by on-column cleavage and removal of the purification tag, leaving only the natural sequence and no affinity tag remnants(*69*). Isotope labeled (^15^N, ^13^C/^15^N, and ^2^H/^13^C/^15^N) samples were prepared using standard procedures(*70, 71*). The GA-prodomain-PXN and GA-prodomain-FAT constructs were transformed into BL21DE3 *E coli* cells and grown in M9 minimal media at 37°C until an OD_600_ of 0.6-0.9 was obtained. Expression was then induced with 1 mM IPTG for 18 h at 25°C. Cells were centrifuged, re-suspended in buffer A (100 mM KPi, 0.1 mM EDTA, pH 7.0), and lysed using sonication. After further centrifugation, the soluble fraction was loaded onto an immobilized subtilisin column (Potomac Affinity Proteins), and washed successively with 5 column volumes of buffer A, 20 column volumes of buffer B (100 mM KPi, 500 mM NaCl, 0.1 mM EDTA, pH 7.0), and 5 column volumes of buffer A. The target protein was then cleaved from the purification tag and eluted from the column using buffer C (100 mM KPi, 0.1 mM EDTA, 2 mM imidazole, pH 7.0). Fractions containing at least 95% pure protein, as determined by SDS-PAGE analysis and MALDI, were pooled and concentrated for further analysis. Mutants were made utilizing the Q5 site-directed mutagenesis kit (New England Biolabs) and expressed and purified as described above.

### NMR spectroscopy

NMR samples of different length PXNs and FAT were prepared at concentrations of 150-300 μM in 100 mM potassium phosphate buffer, 1 mM TCEP, 0.1 mM EDTA, pH 7.0. All NMR spectra were acquired on Bruker Avance III 600 MHz and 900 MHz spectrometers equipped with a Z-gradient ^1^H/^13^C/^15^N cryoprobe at 10°C. Backbone resonance assignments were made using three-dimensional HNCACB, CBCACO(NH), HN(CO)CACB, HNCO, HN(CA)CO, and (H)N(CA)NNH experiments. Spectra were processed with NMRPipe(*72*) and analyzed with Sparky(*73*). Secondary ΔCα shifts were calculated from experimental and sequence corrected random coil chemical shifts(*74*). Heteronuclear {^1^H}-^15^N steady state NOEs were measured using a standard pulse scheme with a relaxation delay of 5 s(*75*).

NMR titration experiments between 100 μM ^15^N-PXN N-domain and unlabeled FAT were performed by acquiring two-dimensional ^1^H-^15^N HSQC spectra as a function of increasing FAT concentrations (10, 20, 40, 60, 80, 100, 150, 200, and 300 μM). As binding mostly resulted in peak broadening at the interfacial regions, peak intensity decay proved to be the most reliable indicator of binding affinity. Relative binding constants for individual LD motifs were obtained by measuring *I*/*I*_0_ for each main chain amide in the relevant LD motif, where *I* is the peak intensity at a given concentration and *I*_0_ is the peak intensity in the absence of any added ligand. The fraction bound (1-*I*/*I*_0_) was then plotted against the total concentration of added FAT and curve fitted(*71*). The binding constants reported are an average of the values for each individual residue in the LD motif ± 1SD. Competition binding experiments were carried out by recording 2D ^1^H-^15^N HSQC spectra of the ^15^N-labeled PXN complex with unlabeled FAT as a function of increasing concentrations of the competing unlabeled peptide.

Intermolecular PRE experiments between natural abundance samples of 100 mM PXN and 100 mM ^15^N-FAT were carried out by first reacting the appropriate PXN Cys mutant with 10 molar equivalents of MTSL (Santa Cruz Biotechnology) for 1-2 h at room temperature. Reactions were monitored to completion utilizing MALDI mass spectrometry, with excess MTSL removed by dialysis. The PXN-MTSL was added to 1 molar equivalent of ^15^N-FAT, and a two-dimensional ^1^H-^15^N HSQC spectrum (400 transients, 128 *t*_1_ increments) was acquired on the paramagnetic sample. A second ^1^H-^15^N HSQC spectrum was acquired with matching parameters after reduction with 20 molar equivalents of sodium ascorbate. Peak intensities for the oxidized and reduced states, *I*_ox_ and *I*_red_, were determined with SPARKY. The reciprocal intermolecular PRE experiments between ^15^N-labeled PXN N-Domain and site-specifically MTSL-labeled FAT were performed in a similar manner.

### SAXS measurements

Solution X-ray scattering data were recorded with a Xenocs Ganesha instrument. Copper Kα incident radiation with a wavelength of 1.542Å was produced by the Rigaku MicroMax 007HF rotating anode generator and collimated via 2 sets of scatter-less slits. Scattered radiation was registered with the Pilatus 300K area detector and the transmitted intensity was monitored via a pin diode. Samples were kept at 25°C and exposed to incident X-ray radiation for 32 sequential 900-second frames. Pixel intensity outliers due to background radiation were removed and the 2D data were corrected for detector sensitivity and solid angle projection per pixel. The data were converted to one-dimensional scattering intensity curves, frame-averaged and buffer-subtracted. Buffer-subtracted data sets acquired at sample-detector distances of 1035 mm and 355 mm were merged to extend the angular resolution range. Radii of gyration were extracted via Guinier fits of the lowest angle scattering data.

### Modeling initial PXN bound FAT structures

Hoellerer et al. (*31*) reported several crystal structures with LD2 (PDB ID: 1OW8) and LD4 (PDB ID: 1OW7) bound to FAT. These structures are with different crystal subunits where the same LD helix is bound to opposite FAT faces. For example, in 1OW7, LD4 is bound to either α14 (chain A) or the α23 face (chains B and C). Likewise, in 1OW8, LD2 is bound to either α14 (chain A) or α23 (chain C) faces.

The above structures were used as templates for homology modeling of the initial PXN/FAT configurations, followed by AWSEM coarse-grain MD simulations to collapse the PXN IDR domains around FAT. For the LD2 and LD4 bound models, both LD2 and LD4 helices were modeled based on the crystal structures. For the LD1 and LD2 bound models, LD2 was modeled based on the bound LD2 of the crystal structure (PDB ID: 1OW8), while LD1 was modeled using the crystal structure of LD4/FAT as template (PDB ID: 1OW7). In all models, the disordered PXN regions were modeled as random loops and LD1, LD2 and LD4 were modeled as helices, based on our NMR results. The following templates were used for the four models: LD1-14, LD2-23 – 1OW8: chain C, 1OW7: chain A; LD1-23, LD2-14 – 1OW8: chain A, 1OW7: chain B; LD2-14, LD4-23 – 1OW8: chain A, 1OW7: chain B; LD2-23, LD4-14 – 1OW8: chain C, 1OW7: chain A. The homology models were generated using MODELLER version 9, using the loopmodel module(*76*). For each PXN/FAT configuration, 50 alternative models were generated, followed by the selection of the best model by the DOPE score(*77*). These structures were then subjected to AWSEM coarse-grain simulations prior to atomistic MD, as described next.

### AWSEM coarse-grain MD simulations of PXN-FAT complex

Each PXN/FAT structure was prepared using the PdbCoords2Lammps.sh script obtained from the AWSEM Github repository (https://github.com/adavtyan/awsemmd). Following the guidelines for simulating IDPs, the helical energy parameter in *fix_backbone_coeff.data* was reduced from 1.5 to 1.2(*32*). Additionally, FAT, LD1, LD2 and LD4 were treated as rigid bodies. The integration timestep was set to 2 femtoseconds. All simulations were performed in the NVT ensemble using nonperiodic boundary conditions and the Nose-Hoover thermostat for temperature control. The systems were initially minimized for 100,000 steps using the Hessian-free truncated Newton method in LAMMPS(*78*), followed by temperature annealing. During this step, starting from randomly assigned velocities at 600K, the temperature was gradually reduced to 300K over 500,000 steps. The last frame from each MD trajectory was converted into atomistic backbone coordinates using the *BuildAllAtomsFromLammps.py* script from AWSEM.

The fully atomistic structures of the PXN-FAT complex including residue side-chains (to be used as inputs for the atomistic MD) were reconstructed from the backbone coordinates from AWSEM using homology modeling in MODELLER. FAT was modeled based on the crystal structure coordinates (PDB IDs: 1OW7, 1OW8), while PXN was modeled based on the backbone coordinates from AWSEM. The final structure for each PXN orientation was selected from a pool of 50 structures based on lowest DOPE score.

### System setup for atomistic MD

Resulting structures from the AWSEM simulations were subjected to thorough minimization using the PrimeX module of Schrodinger(*79*), followed by addition of hydrogen atoms using the pdb2gmx module of GROMACS 5.1(*80*). Each structure was solvated in explicit water in a cubic box with a separation of 25Å maintained between the protein atoms and the simulation box boundaries. Sodium and chloride ions were added to neutralize the charges and create an effective ion concentration of 0.15 mM. The total number of atoms in the four systems were between 280,000 to 450,000. The protein atoms and water molecules were parameterized using the Amber99SB-disp force-field(*81*) and the TIP4P-D water model respectively(*82*), designed to simulate both folded and disordered protein conformations. The prepared systems were converted to the AMBER format using the gromber module of ParmEd(*83*). Hydrogen mass repartitioning was applied to the protein hydrogen atoms to enable an integration timestep of 4 fs(*84*). All simulations were performed using the AMBER 18 software package on a GPU cluster.

### System equilibration and conventional MD

Each system was initially minimized in two steps: 1) 20,000 steps of restrained minimization with protein heavy atoms restrained via a harmonic force constant of 5000 kcal/mol; 2) 200,000 steps of fully unrestrained minimization. Next, the systems were heated from 0 to 310K over 30 ns in the NVT ensemble, during which the protein heavy atoms were subjected to harmonic positional restraints (force constant: 500 kcal/mol). Following this, the systems were equilibrated in the NPT ensemble (temperature: 310K, pressure: 1 atm) for 70 ns, during which the heavy atom restraints were gradually reduced to zero. Finally, the fully unrestrained systems were subjected to MD simulations in the NPT ensemble for another 700-1000 ns, using a timestep of 4 fs. Temperature and pressure were maintained using Langevin dynamics scaling with a collision frequency (*gamma_ln*) of 1 and pressure relaxation time of 2 ps.

### Gaussian accelerated MD

Starting from the last frame of the conventional MD of each system, GAMD equilibration was performed using the following parameters: igamd=3 (dual potential and dihedral energy boost), ntcmd = 1000000, nteb = 25000000, ntave = 200000, ntcmdprep = 200000, ntebprep = 800000, sigma0P = 6.0, sigma0D = 6.0. Total equilibration time was 50 ns using an integration timestep of 2 fs. Following the equilibration, five independent production runs were performed starting with random velocities, lasting between 1.2 to 1.4 μs each. A 2 fs integration timestep was used during the GAMD simulations. The simulation snapshots were recorded every 10 ps. In total, we performed 20 independent GAMD simulations (5 x 4 configurations), resulting in an ensemble of 2.8 million structures.

### Calculation of PRE intensity ratios from MD ensembles

The PRE intensity ratios were calculated from the MD derived PXN/FAT structures following the Solomon-Bloembergen equations (*84, 85*). Briefly, the transverse relaxation rates in the presence of the spin label in oxidized 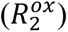 and reduced states 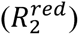 are related by 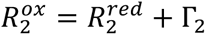 (equation 1), where Γ_2_ is the weighted average PRE enhancement of a given residue from a structural ensemble of N conformations, given by 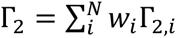(equation 2), with *w_i_* being the weight of the i^th^ frame. To estimate Γ_2,*i*_ for each MD conformation, we used the Solomon-Bloembergen equation:

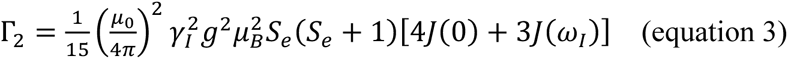

where *ω_I_* and *γ_I_* are the Larmor frequency and gyromagnetic ratio of the proton respectively, *S*_e_ = 1/2 is the electron spin quantum number, *μ_0_* is the free space permeability, *μ_B_* is the Bohr magneton, and *g* is the electron g-factor. The spectral density function *J(ω)* is given by:

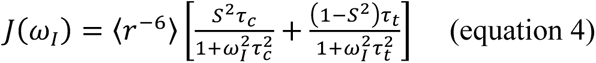

where, *t_c_* is approximately the rotational correlation time of the protein and *t_i_* is the correlation time of the spin label, set to 500 ps; *t_c_* was estimated to be 29 ns based on the molecular weight of the PXN/FAT complex (47.8 kDa)(*86*); *r* is the distance between the unpaired electron of the spin label and backbone amide proton of a given residue; 〈*r*^−6^〉 represents the average over possible rotamer states of the spin label, weighted by their respective Boltzmann probabilities (*35*); *S* is the generalized order parameter given by 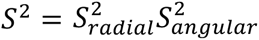 (equation 5), with 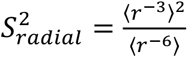 (equation 6). The values of 〈*r*^−3^〉, 〈*r*^−6^〉 and 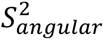 were calculated for each MD frame using the DEER-PREdict method as described(*35*). Utilizing the values of the above parameters, Γ_2_was then calculated. Assuming an exponential decay of the proton magnetization intensity through transverse relaxation within the total INEPT time *t_d_* of HSQC measurement (10 ms), the PRE intensity ratio can be estimated as 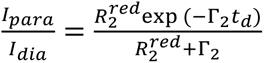 (equation 7). Based on the molecular weight of the PXN/FAT complex and comparison with measured R_2_ from proteins with different molecular weights(*87*), 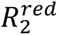 was set to 40s^-1^.

### Reweighting of PXN/FAT MD ensemble using Bayesian Maximum Entropy (BME)

Details of the BME method have been described elsewhere(*33*) and previously applied in reweighting MD derived IDP ensembles(*34, 88–90*). Here we discuss the specifics of the approach as they apply to the current work. Using BME, we refined the MD ensemble by reweighting each MD frame, leading to improved agreement of the weighted average PRE values with experiment. In total, we used 730 experimentally determined PRE intensity ratios at PXN residues from four different spin labels placed on FAT (**Fig. 5C,F,I,L**). To facilitate BME fitting, the experimental PRE intensity profiles per spin label were smoothed using locally estimated scatterplot smoothing(*91*) (LOESS, α = 0.2) (**Fig. S6**). Next, the intensity ratios were converted to Γ_2_ employing equation 7. Notably, due to the exponential relation between intensity ratio and Γ_2_ (**Fig. S7**), small values of intensity ratios can produce astronomically large Γ_2_ values, leading to numerical instability in the BME algorithm. To alleviate this, Γ_2_ values were truncated at 450, beyond which any further increase in Γ_2_ did not appreciably change the intensity ratio.

The BME procedure is described in **Fig. S7**. Given a specific θ and a set of initial weights 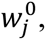 a set of Lagrange multipliers λ_i_ were first derived by minimizing the cost function:

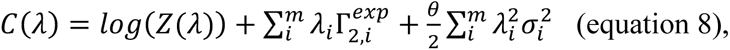

where *m* is the number of experimental Γ_2_values from the four spin labels, θ is an adjustable parameter describing the trade-off between agreement with experiments versus entropy of the weight distribution, 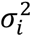 is the experimental error (i.e. variance) associated with each data-point, and Z is a normalization factor to ensure that the weights add up to 1. The weight for each frame *j* can then be derived using the relation:

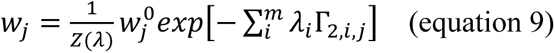

where Γ_2,*i*,*j*_ is the Γ_2_ value for the *j^th^* frame. Starting with uniform initial weights, we tested the BME fitting for a range of θ values. The variance scaled mean square error 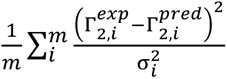 (equation 10) was then plotted as function of θ (**Fig. S7C**). The optimal θ was chosen as the one beyond which χ^2^ changed minimally with decreasing θ. We initially performed the BME calculation by keeping 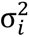 constant, which led to low fitting error at low PRE ratios, but higher error at PRE ratios > 0.6 (**Fig. S8A**). To mitigate this, we let 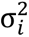 vary inversely with the experimental PRE, leading to a balanced distribution of fitting errors throughout the entire range of PRE ratios (**Fig. S8B**).

### Representing conformations in reduced dimension space

The FAT-bound PXN N-domain conformations were analyzed and clustered in reduced dimension space using Uniform Manifold Approximation and Projection (UMAP) as follows. Briefly, the PXN chain was divided into consecutive 4 amino acid segments and the minimum C_α_ distances between segment pairs were calculated for each MD frame (**Fig. S9A**). To avoid including highly correlated distances, only segment pairs separated by more than two segments in between were considered, leading to 2775 distances. These distances were used to perform a principal component analysis (PCA) (**Fig. S9B**) and, utilizing the top 50 PCs, the UMAP coordinates were calculated. In time series data such as MD, where temporally close conformations tend to be strongly correlated, UMAP can emphasize these correlations rather than true structural similarities learned from the global ensemble. To address this, we first generated the UMAP by training against a small subset of randomly selected conformations (10% of the total ensemble) followed by projection of the rest of the PXN conformations to this space. The PXN conformations were grouped into 96 clusters using the top 50 PC based coordinates, employing the K-nearest neighbor (KNN) similarity measure and the Louvain clustering algorithm(*92*). The segment-pairwise distances were calculated using the MDanalysis package(*93*). The UMAP calculation and clustering were performed using the uwot(*94*) and Seurat(*95*) R packages respectively. The analysis pipeline was implemented in R 4.3(*96*).

To assess the validity of the UMAP representation, we reasoned that a successful UMAP transformation should capture the similarities and differences among the different FAT-bound PXN conformations, thereby recovering the clustering structure. We therefore clustered the PXN conformations using their UMAP coordinates employing the K-means algorithm and compared the resulting clustering assignments to those obtained using PC based Louvain clustering. Positive adjusted Rand Index (ARI) and adjusted Mutual Information (AMI) between the two clustering assignments indicated strong correspondence between the UMAP based and PC based clusters (**Fig. S9C**). We further estimated the statistical significance of the ARI and AMI measures by randomly shuffling the UMAP based cluster labels among PXN conformations 10,000 times, generating null distributions (**Fig. S9D & E**). The null ARI and AMI values from the randomly generated clusters were close to zero (ARI: -2.5×10^-7^±5.5×10^-6^, AMI: -2.7×10^-6^±5.4×10^-5^ (mean±SD)), indicating strong statistical significance.

### MD simulation of unbound PXN N-domain

PXN N-domain structures were modeled as a random coil using Modeller (*76*), except LD1, LD2 and LD4, which were modeled as α-helices. The modeled PXN structure was subjected to coarse-grained MD using AWSEM (Atomistic associative memory Water-mediated Structure and Energy Model)(*32*). The system was initially subjected to minimization and annealing using the same protocol as the PXN-FAT complex simulations. Next, an equilibration was performed at 350K in the NVT ensemble for 50,000,000 steps. The production runs consisted of five independent simulations at 350K for 1.25×10^9^ steps, starting with different random velocities, leading to 750,000 conformations. During coarse-grained MD, LD1, LD2 and LD4 were treated as rigid bodies and the helical energy parameter in *fix_backbone_coeff.data* was reduced from 1.5 to 1.2. Using the default parameters, AWSEM produced highly compact PXN structures with average *R*_g_ ∼ 20Å. To sample more realistic conformations with larger *R*_g_, the desolvation barrier V_DSB_ (first line of the section “Solvent_Barrier” in *fix_backbone_coeff.data*) was increased from its default value of zero. We tested several values of V_DSB_, finally settling for V_DSB_ = 1.0, which gave an average *R*_g_ (56Å) in agreement with experiment. The ensemble of protein backbone conformations from AWSEM were converted to all-atom structures by adding side-chains using the program SCWRL(*97*). To add side-chains, each backbone conformation from AWSEM was individually exported as PDB and side-chains were added using the SCWRL(*97*). The output PDB files were then concatenated into a multi-frame PDB and converted into DCD using CatDCD(*98*).

### Structure visualization and analysis

The PXN/FAT complex and unbound PXN trajectories were visualized using VMD(*99*) and PyMOL(*100*). Radius of gyration was calculated using the CPPTRAJ module of AMBER 18(*101*).

### Multiple sequence alignment of PXN

The PXN sequence was used to search the nonredundant protein database in Uniprot(*102*) for homologous sequences. Isoform hits with documented protein-level evidence were selected for alignment using Clustal-Omega(*103, 104*). The alignment figure was prepared using JalView(*105*).

### Configurational entropy along PXN sequence

Entropy in macromolecules is contributed by several aspects including solvent degrees of freedom as well as solute configurational entropy consisting of vibrational motion, protein center of mass movement and collective motion arising from backbone and sidechain dynamics(*106*). The configurational entropy can be further subdivided into vibrational and conformational entropy (entropy associated with the number of discrete conformations sampled by the macromolecule)(*107*). We calculated the conformational entropy per PXN residue in all four FAT bound configurations by employing an information-theoretic approach (*40, 41*). Detailed description of the entropy calculation is given elsewhere(*40*). Briefly, the backbone (excluding the ω angle, which has limited flexibility) and sidechain torsion angles of every amino acid were computed for each frame of the MD trajectories (**Fig. 10A**). Next, the torsion angle ranges were divided into 35 bins and the BME reweighted frequency distribution was determined for each torsion angle in the four FAT bound configurations (**Fig. 10B**). The Shannon entropy for a given torsion angle was calculated using a modified version of the following relation(*41*):

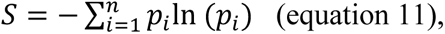

where *p_i_* is the BME reweighted probability of the *i^th^*bin and *n* is the total number of bins. After applying corrections for using discrete bins and undersampling, the modified expression for entropy becomes(*108, 109*):

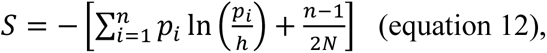

where *h* is bin-width and *N* is the number of frames in the trajectory. Entropy values for all torsion angles belonging to a given residue were summed to calculate residue-wise entropy. Entropy calculations were performed using the MDAnalysis package(*93*) and Python 3.8(*110*).

## Author contributions

Conceptualization: SB, RS, JO, PK, YH

Methodology: SB, JO, AG, YH, AM

Investigation: SB, AG, JO, YH, YC

Resources: AG, JO, SB, RS, AM

Data curation: AG, JO, SB

Validation: AG, YC, PK, JO, SB

Formal analysis: AG, YC, JO, SB

Software: AG, SB

Visualization: SB, JO, AG

Project administration: PK, SB, RS, JO

Supervision: SB, JO, RS

Funding acquisition: JO, RS

Writing—original draft: JO, PK, SB, AG

Writing—review & editing: SB, AM, PK, JO, AG, RS

## Data and materials availability

Backbone NMR resonance assignments for H_N_, N, Cα, Cβ, and CO were deposited in the Biological Magnetic Resonance Data Bank (BMRB) with the following accession codes for PXN LD1-2 (51553), LD2-4 (51554), N-domain (51555), and FAT (51556). The PRE profiles, initial PXN/FAT models, downsampled MD trajectories, and analysis/figure scripts are deposited at Zenodo with DOI:10.5281/zenodo.13836208. All data needed to evaluate the conclusions in the manuscript are present in the main text and/or the Supplementary Materials.

## SUPPLEMENTARY MATERIALS

**Figure S1:**
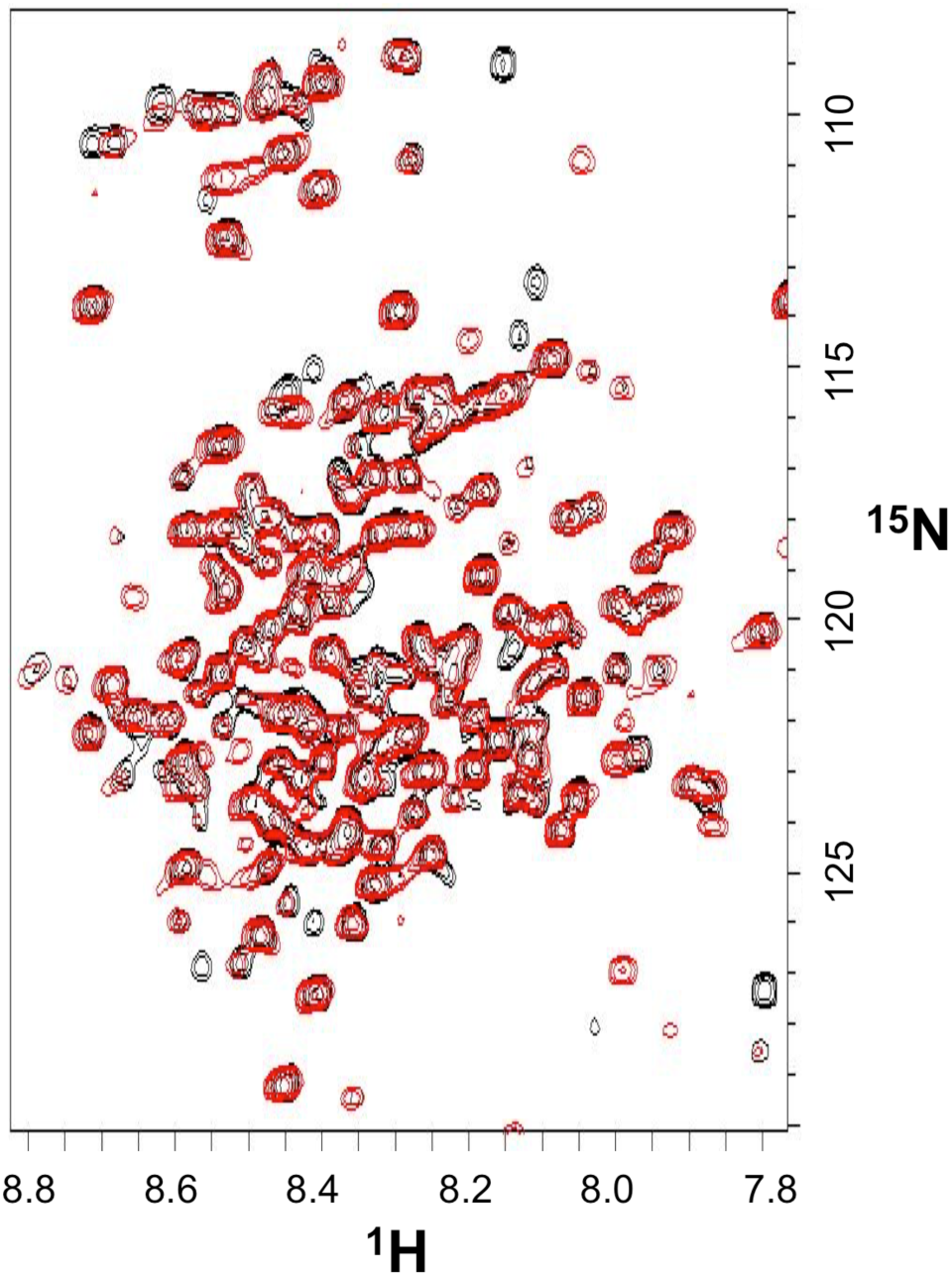
Comparison of full-length PXN and N-domain PXN NMR spectra. Overlaid two dimensional ^1^H-^15^N HSQC spectra of the 311-residue N-domain of PXN (black) and 557-residue, full-length PXN (red). Peaks due to the ordered LIM domains in full-length PXN are broadened and not readily apparent, presumably due to their slower tumbling relative to the more flexible N-domain.

**Figure S2:**
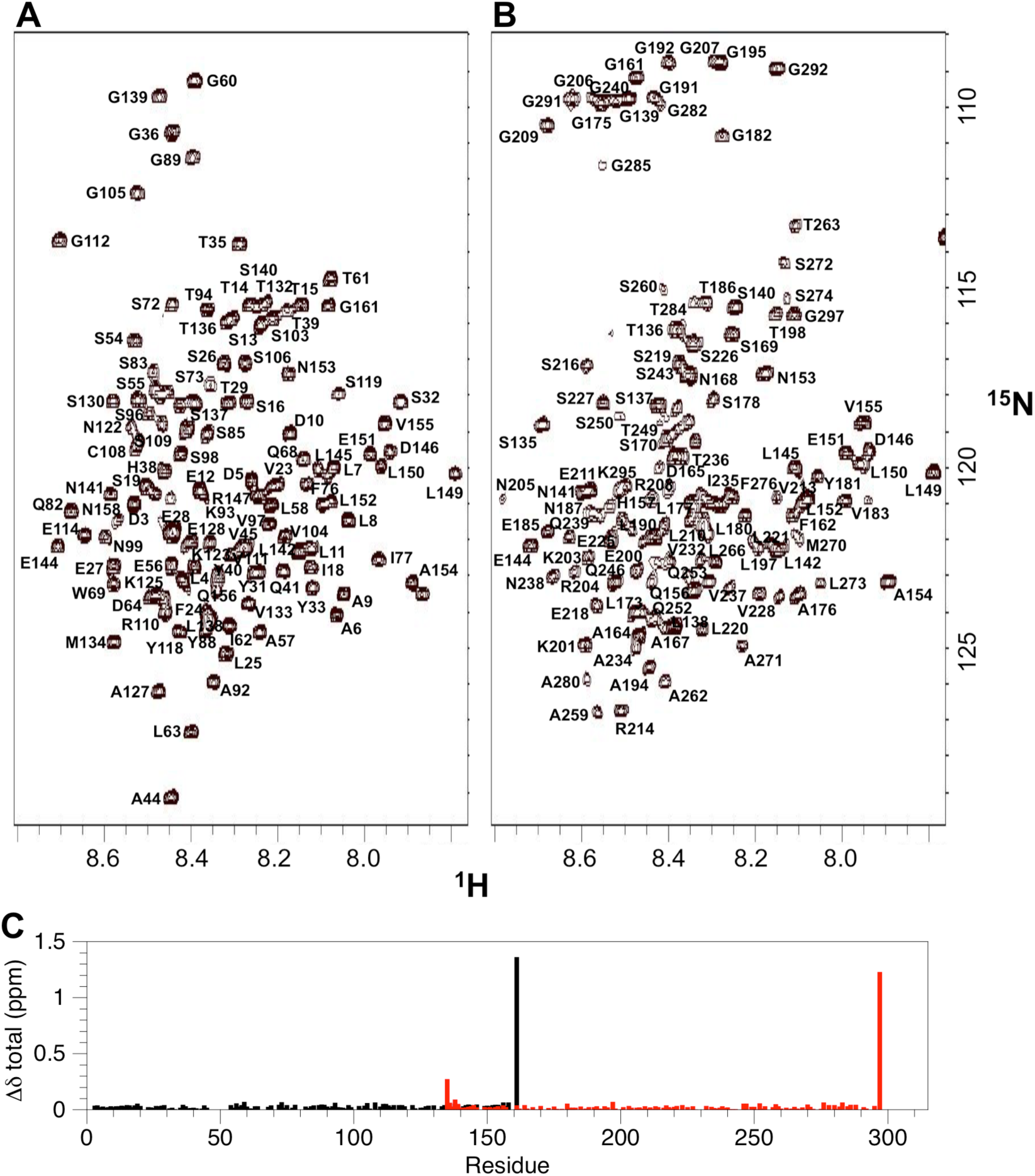
Backbone amide assignment of PXN fragments. Two dimensional ^1^H-^15^N HSQC spectra for (**A**) PXN LD1-2 and (**B**) PXN LD2-4 with backbone amide chemical shift assignments. (**C**) Backbone amide chemical shift perturbations between the PXN N-domain (LD1-5) and the corresponding residues in LD1-2 (black) and LD2-4 (red). The chemical shift perturbations were determined using Δ8_total_ = [(*W*_H_Δ8_H_)^2^ + (*W*_N_Δ8_N_)^2^]^1/2^, where *W*_H_ = 1 and *W*_N_ = 0.2

**Figure S3:**
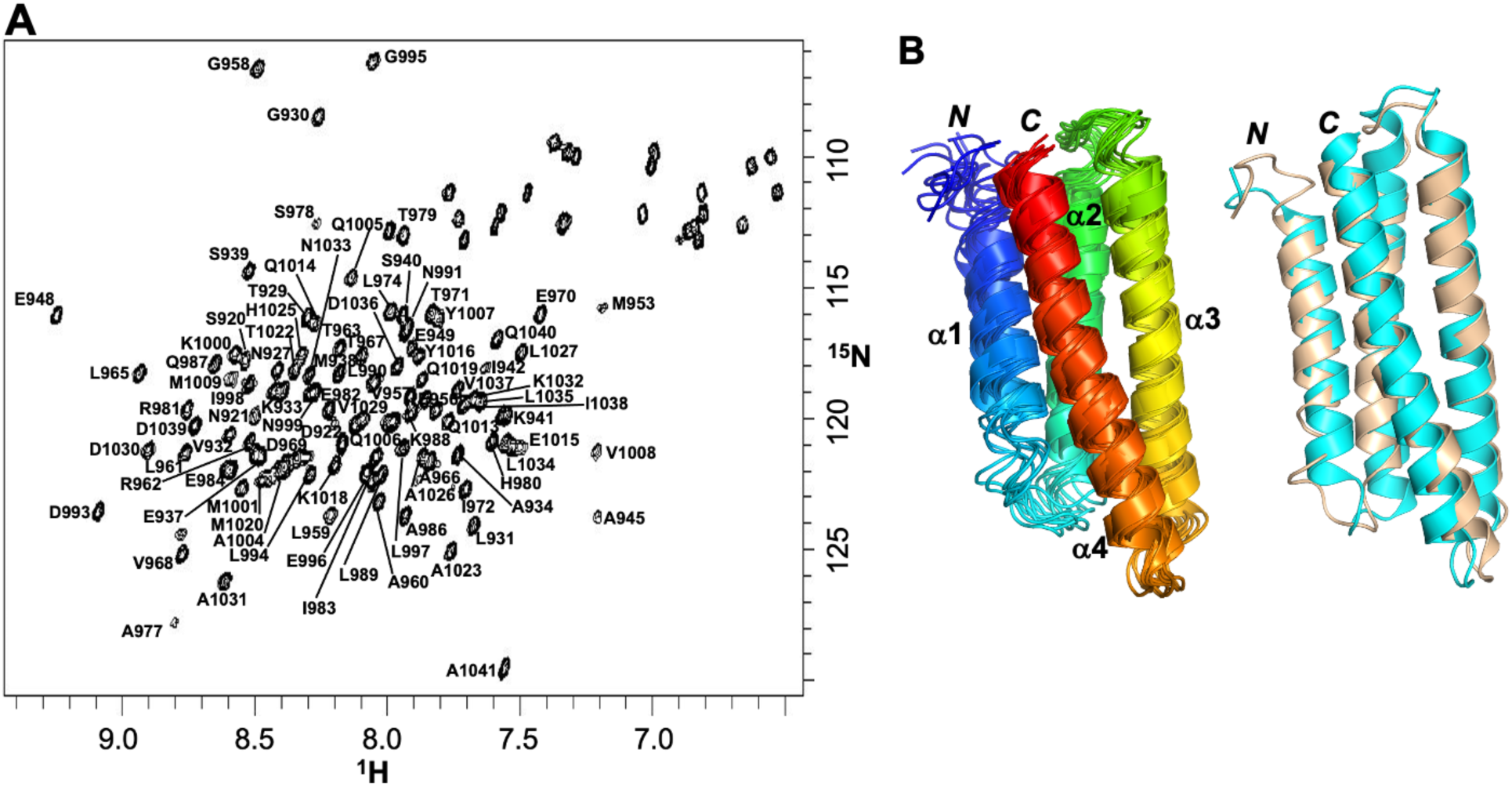
NMR structural analysis of human FAT. Structural analysis of human FAT. (**A**) Two dimensional ^1^H-^15^N HSQC spectrum of human FAT with backbone amide peak assignments. (**B**) CSRosetta structure showing the ensemble for the 10 lowest energy conformations (left), deposited in PDBDev (Accession code 00000391). Superposition with the X-ray structure (PDB 1OW8, cyan) gives a backbone RMSD of 1.5 Å (right). See Table S1 for structure statistics.

**Figure S4:**
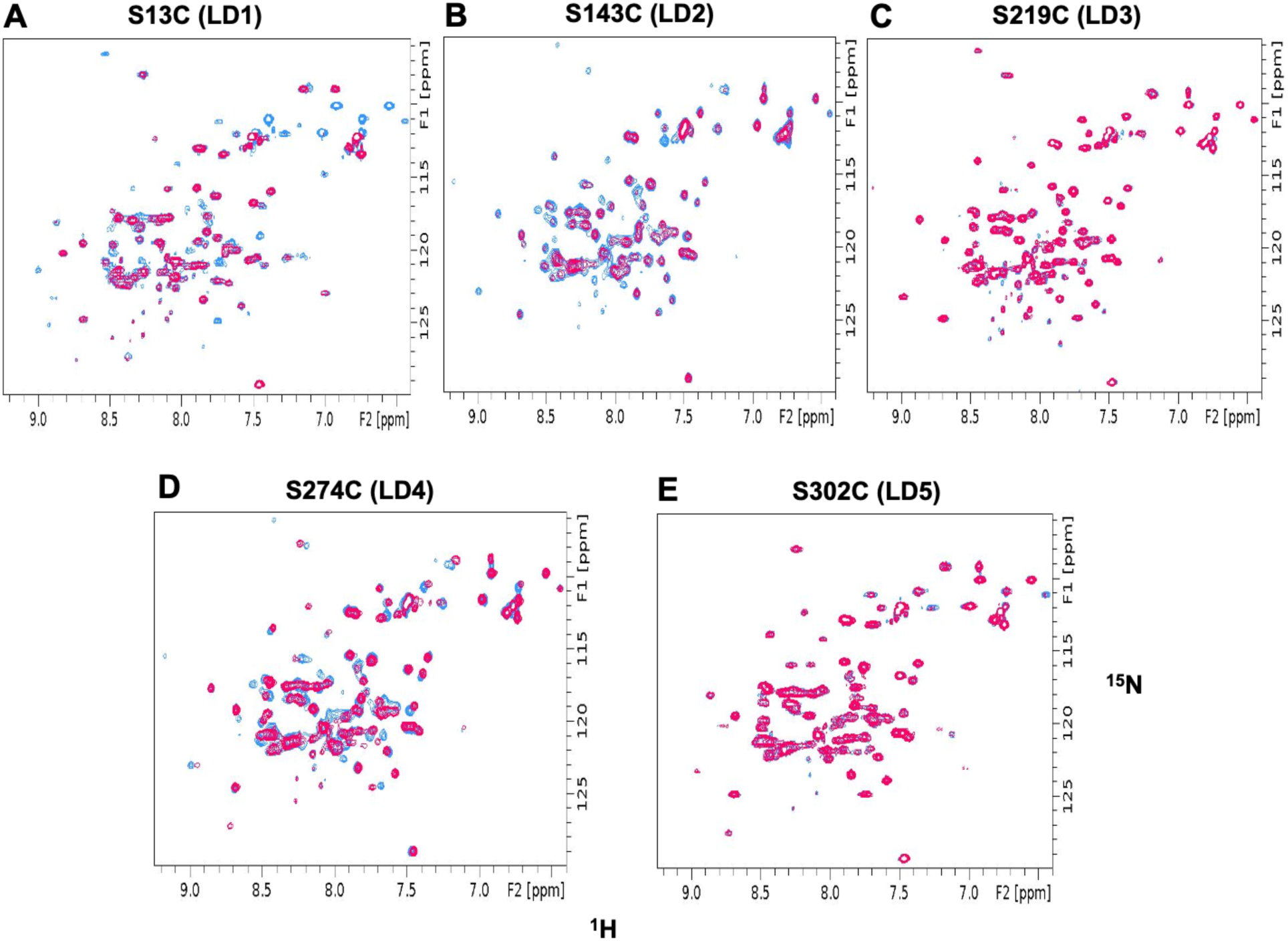
Intermolecular PRE data for determining how PXN binds FAT. Two dimensional ^1^H-^15^N HSQC spectra of the FAT domain used for mapping binding epitopes of LD motifs onto the FAT surface, as described in Figure 4. Overlaid spectra for reduced (blue) and oxidized (red) states are shown for each PXN MTSL-spin label position as indicated.

**Figure S5:**
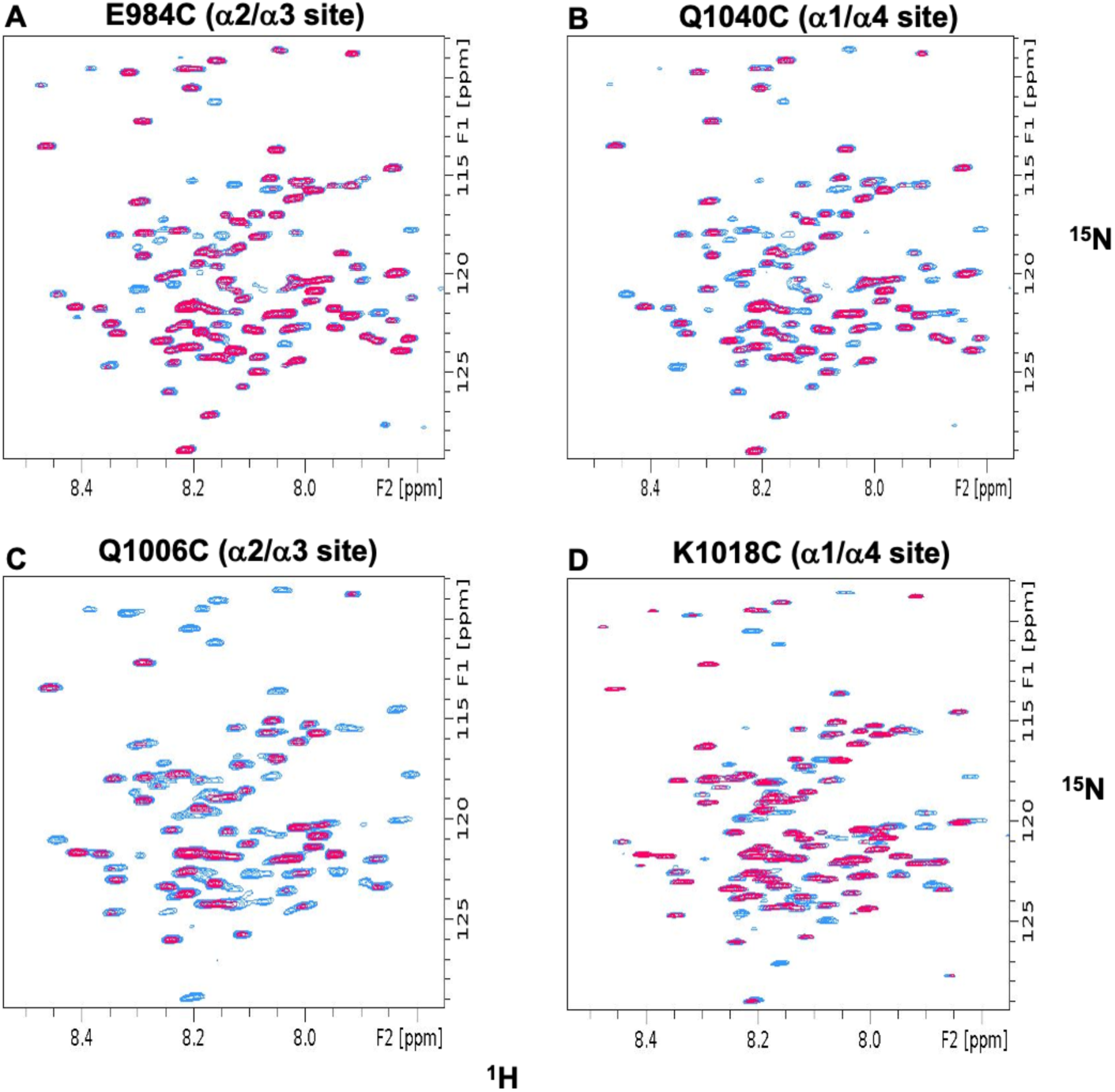
Intermolecular PRE data probing flexibility of the FAT-bound PXN chain. Two dimensional ^1^H-^15^N HSQC spectra of the PXN N-domain used to probe its conformational dynamics around the α2/α3 and α1/α4 sites of the FAT domain, as described in Figure 5. Overlaid spectra for reduced (blue) and oxidized (red) states are shown for each FAT MTSL-spin label position as indicated.

**Figure S6:**
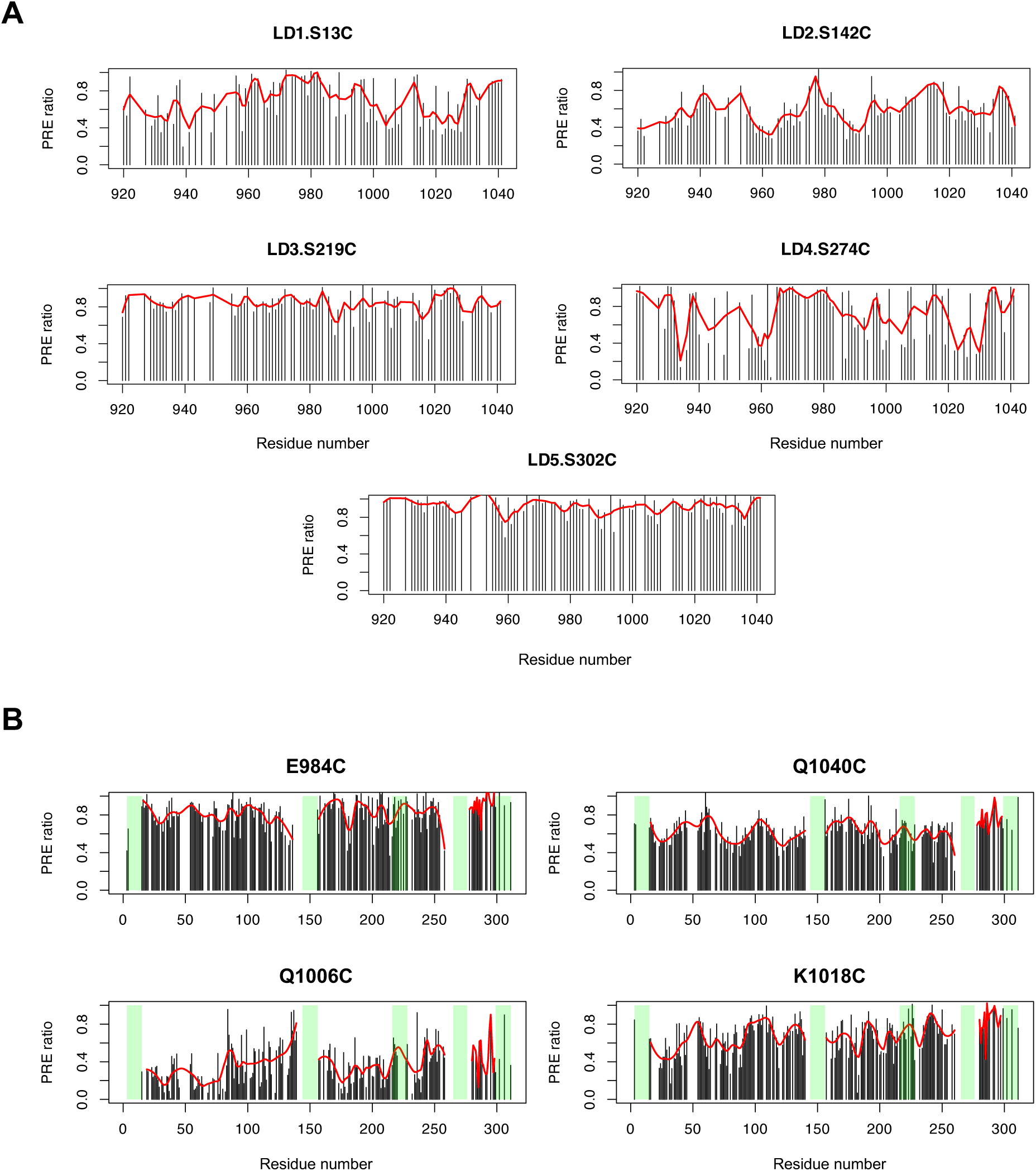
Comparison of smoothed and unsmoothed intermolecular PRE profiles. Smoothing of the experimental PRE profiles using the LOESS algorithm for MTSL probes along (**A**) PXN sequence, (**B**) FAT sequence. Black lines represent the original data and the red curves the smoothed profiles. For each plot in panel B, the positions of the LD motifs are highlighted in green.

**Figure S7:**
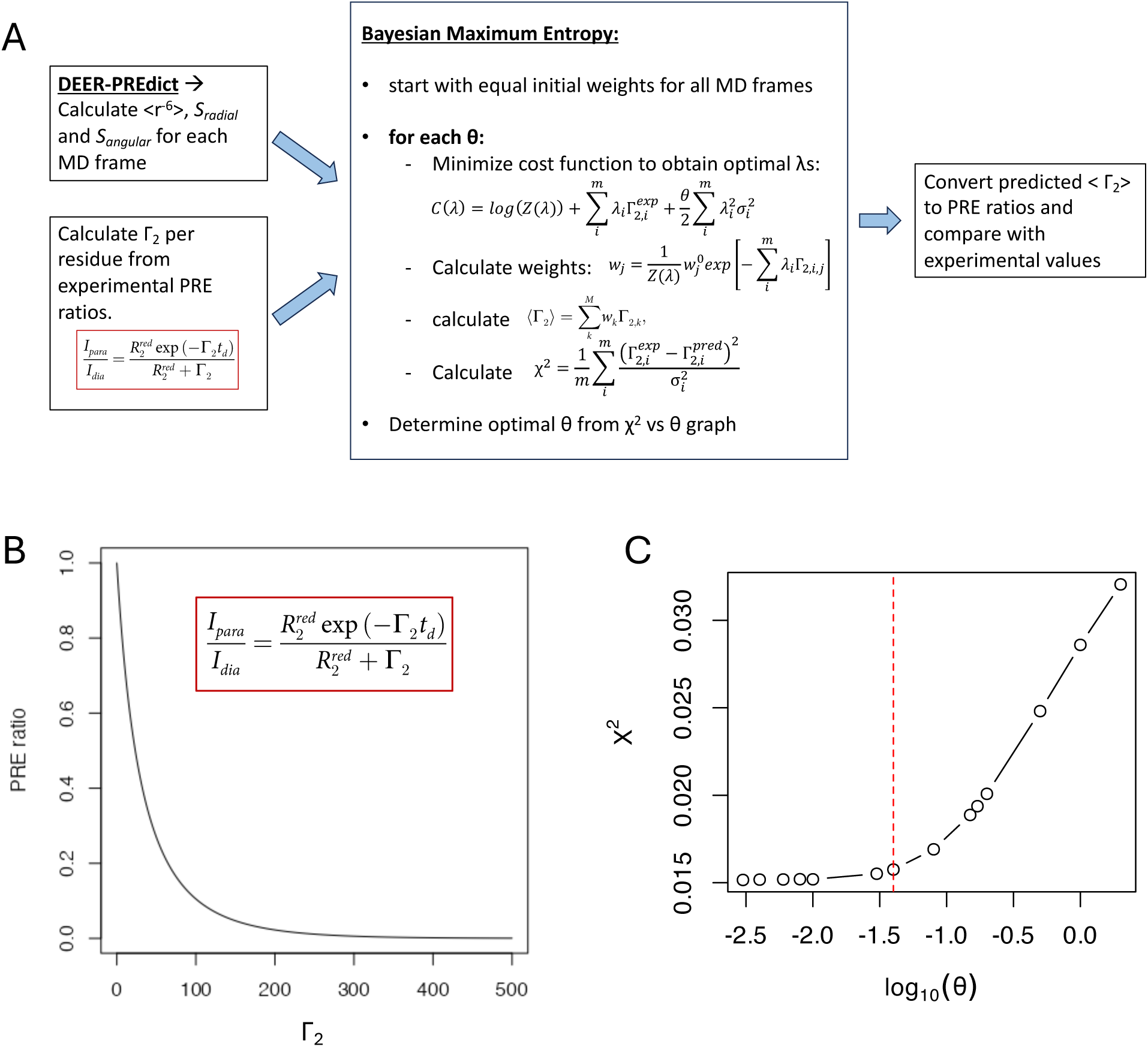
Computational pipeline for deriving trajectory frame weights using experimental PRE ratios. (**A**) Schematic describing the iterative BME procedure for deriving trajectory weights in conjunction with experimental PRE ratios. (**B**) Relationship between ρ_2_ and PRE intensity ratio, as reflected by the equation in the red box. (**C**) Variance scaled mean square error χ^2^as function of θ. The optimal θ is marked by the vertical red dashed line.

**Figure S8:**
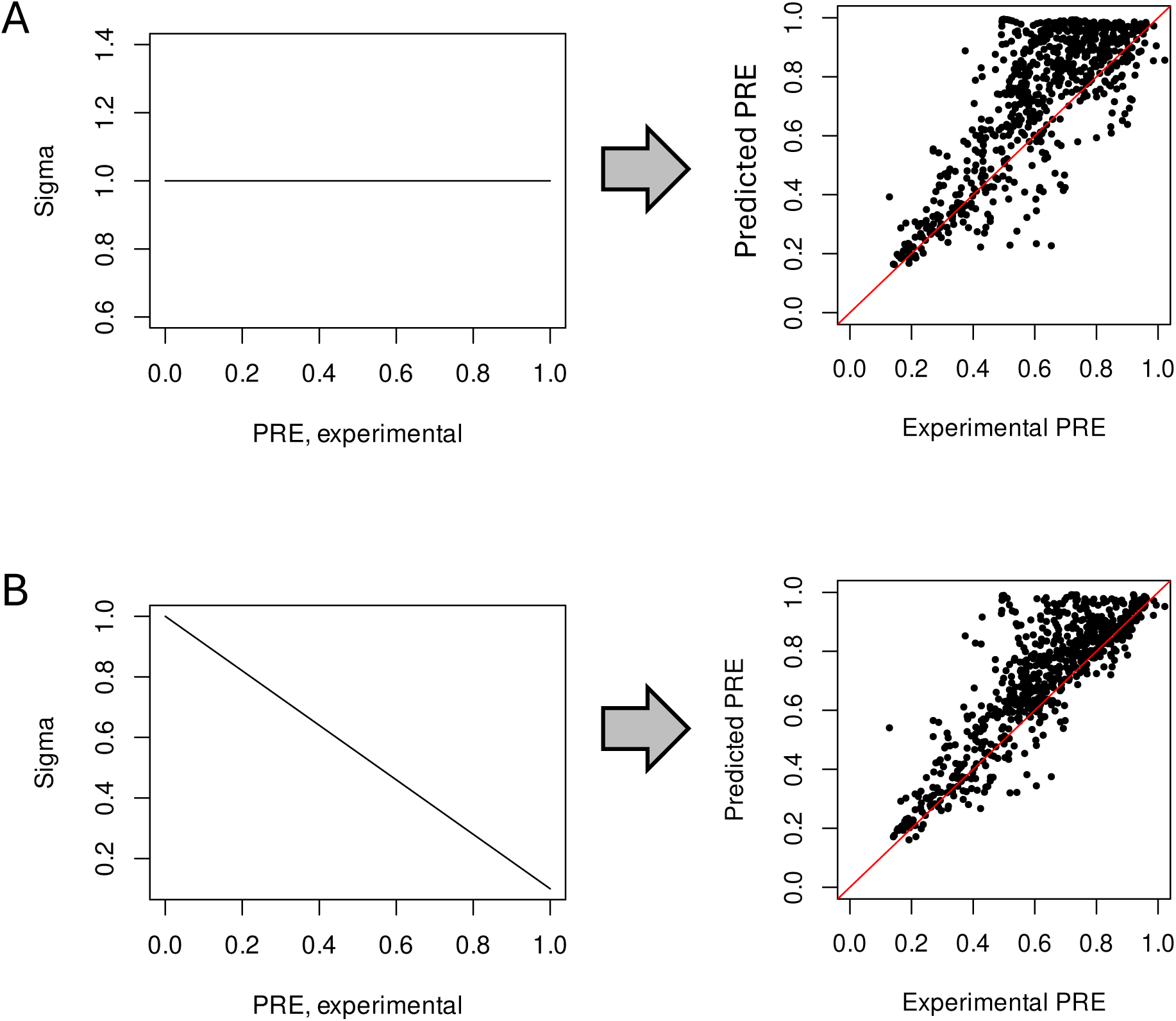
Applying variable sigma in deriving trajectory frame weights using the BME method. Comparison of PRE correlations obtained using constant (**A**) versus variable sigma (**B**) in the BME equation (see Methods).

**Figure S9:**
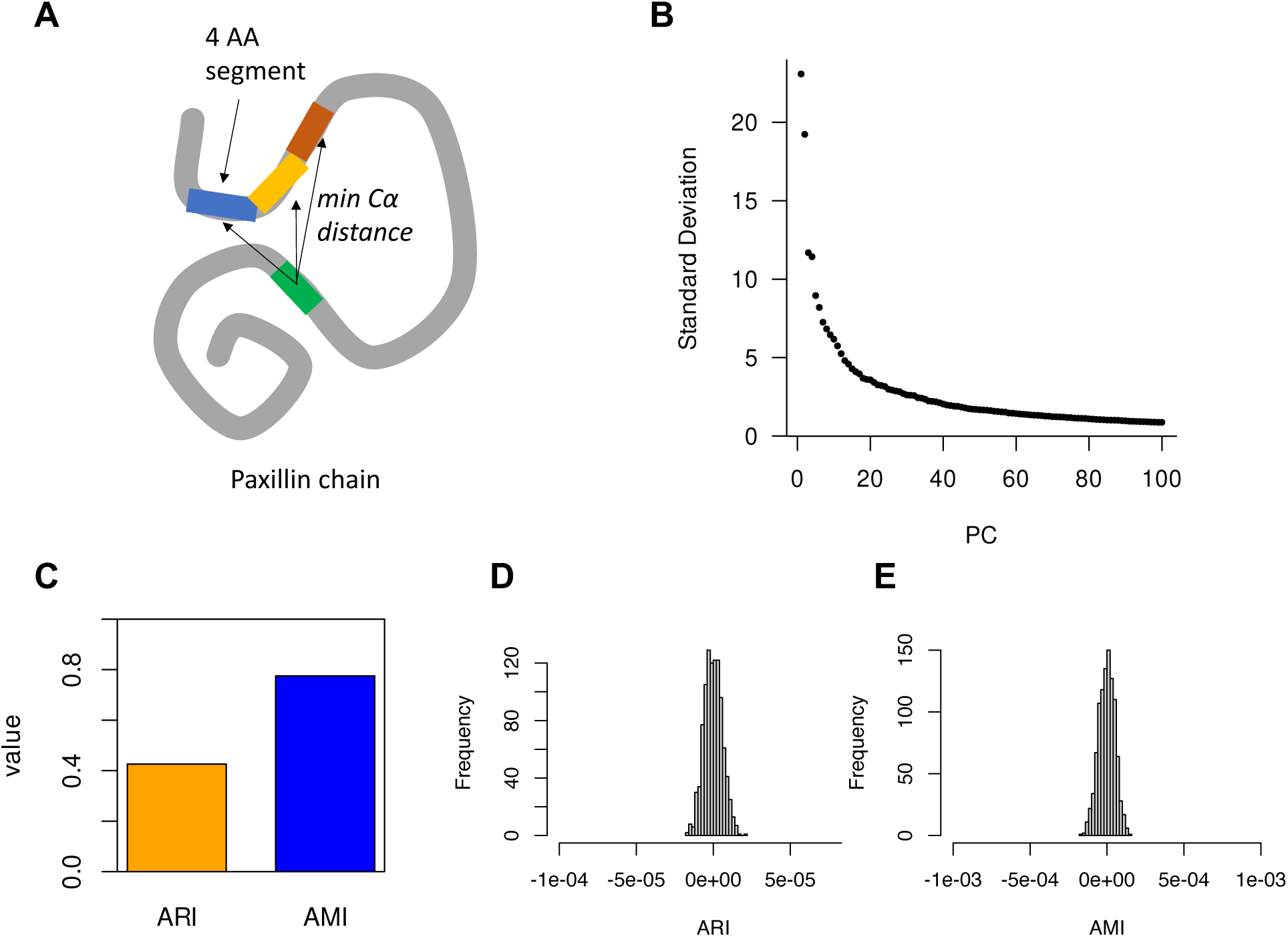
Structure-based similarity metric for clustering PXN MD conformations and UMAP projection. (**A**) Schematic describing the amino acid segments along the PXN chain used in calculating inter-segment distances for deriving the UMAP. (**B**) Percentage of variation explained by the top 100 principal components (PCs) as calculated from the inter-segment distances among the trajectory frames. (**C**) Adjusted Rand Index (ARI) and Adjusted Mutual Information (AMI) between the conformation clusters obtained using the top PCs versus the two UMAP coordinates. (**D-E**) Null bootstrapped ARI and AMI distributions obtained by randomly scrambling the UMAP-based cluster labels 10,000 times and comparing with the PC-based cluster labels.

**Figure S10:**
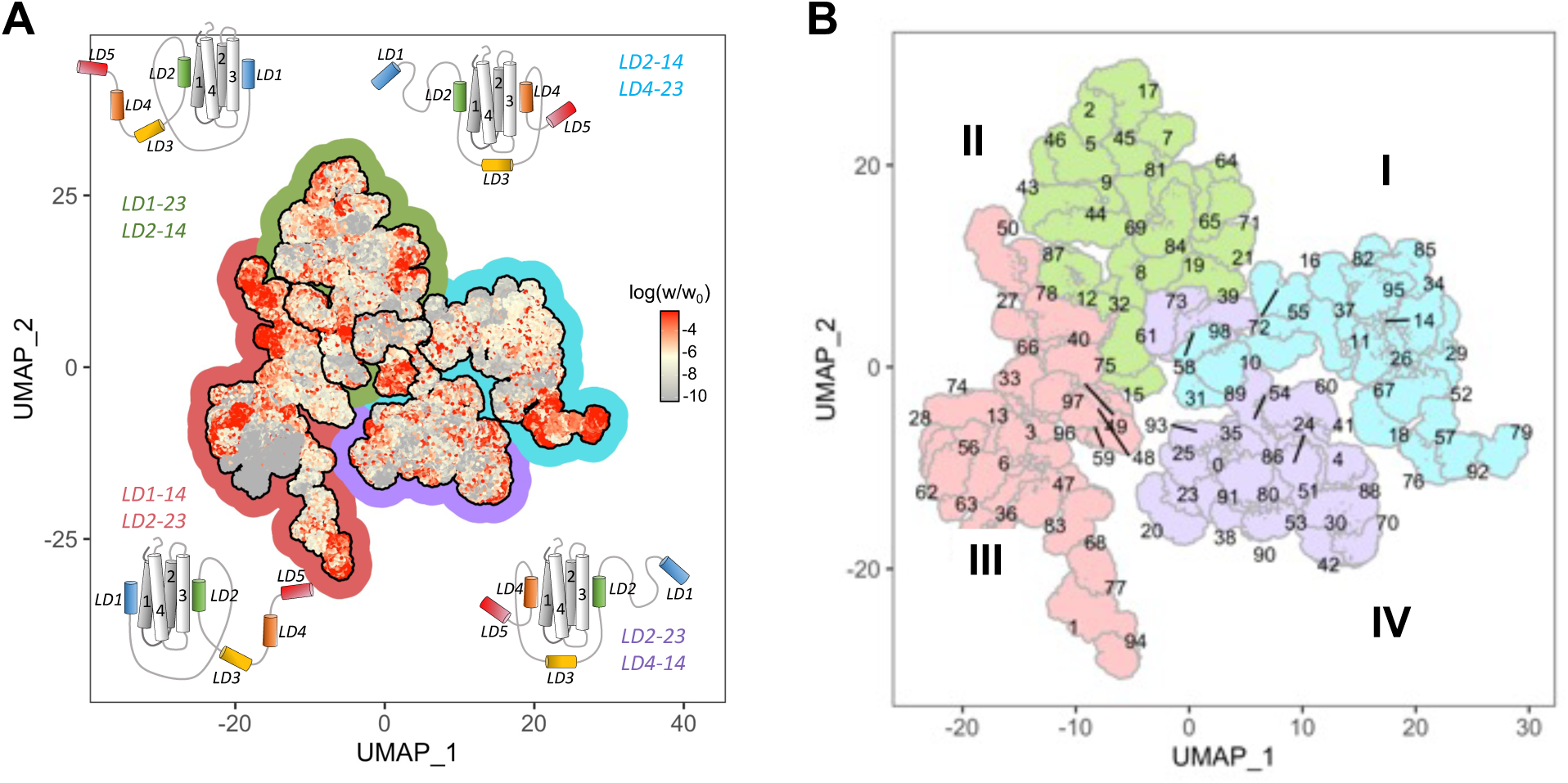
UMAP representation of PXN conformational ensemble. (**A**) MD-generated PXN/FAT ensemble projected in UMAP space, where the UMAP coordinates are derived using PXN inter-chain contacts in each MD conformation (see Methods for details). The conformations are color-coded according to their weights obtained using the BME approach. Red regions in the UMAP contribute more towards the PRE agreement compared to gray regions. Colored halos around each UMAP region are indicative of the PXN/FAT orientation used in the MD simulations. (**B**) MD-derived PXN conformations shown in a UMAP plot. Clusters are labeled using cluster indices, gray lines demarcate cluster boundaries, and conformations are color-coded according to the PXN orientation (I-IV).

**Figure S11:**
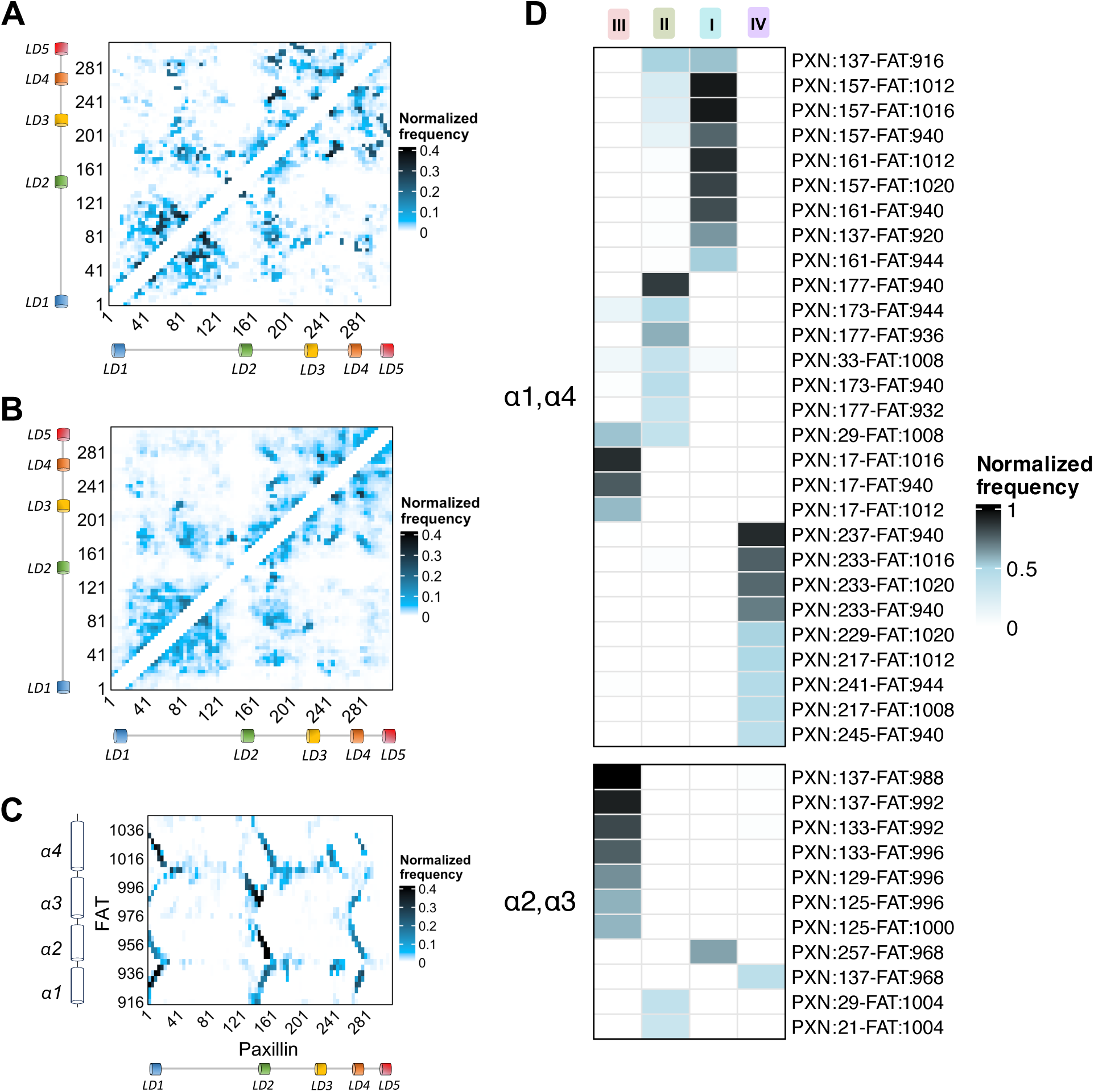
PXN-FAT interaction frequency map from reweighted MD ensemble. (**A**) Contact map of the FAT-bound PXN N-domain conformational ensemble obtained from MD using the BME reweighted ensembles. The PXN N-domain was divided into consecutive 4 amino acid-long segments and the minimum Cα distances between segment pairs were calculated for each MD frame. Two segments were defined to be in contact if their inter-segment distance was less than 8Å (for details, see Methods and Fig. S9A). Cells in the contact map are colored according to the contact frequencies of segment pairs. Short range contacts between consecutive segments were omitted. (**B**) Contact map derived from the original unweighted MD ensemble. (**C**) PXN/FAT contact map using the BME reweighted PXN ensemble. The contact definition is the same as for panels A-B. (**D**) Heatmap depicting the top PXN/FAT linker region contacts per MD-derived state, shown separately for the α1/α4 and α2/α3 faces of the FAT domain.

**Figure S12:**
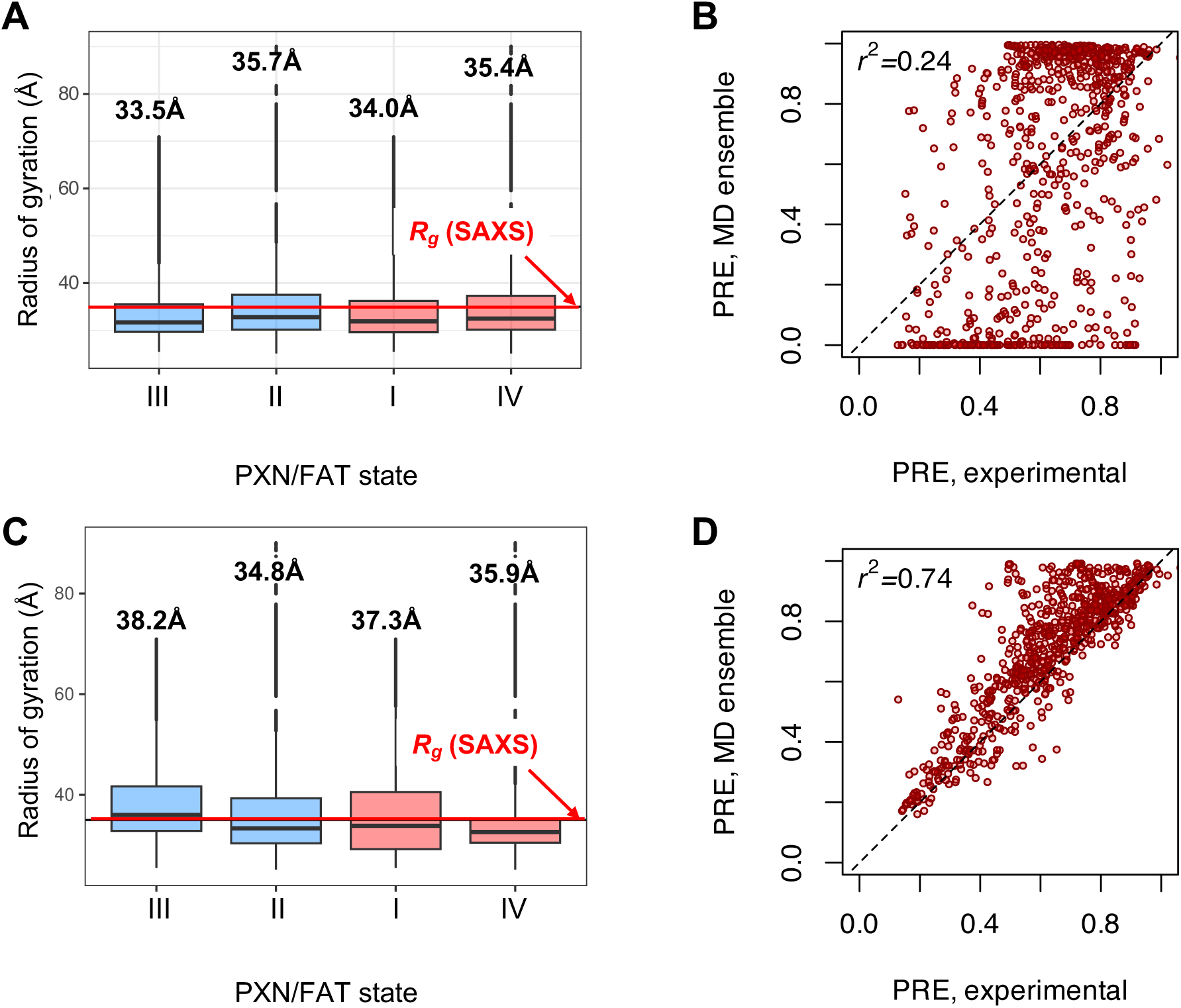
Radii of gyration and PRE correlations from PXN MD ensembles. Radii of gyration and PRE agreement comparison between the original unweighted **(A,B)** and BME reweighted **(C,D)** ensembles, for each of the four PXN/FAT states. **(A, C)** Box and whisker plots showing the radii of gyration (*R*_g_) calculated from the original and BME reweighted ensembles respectively. Boxes represent the interquartile ranges, with the outlier MD conformations plotted along the vertical lines above and below each box. The horizontal red line is the experimental *R*_g_ (35Å) obtained from SAXS. **(B,D)** PRE intensity ratios from all four MTSL probes are compared between their experimental and MD derived counterparts, for the original **(B)** and reweighted **(D)** ensembles. Pearson’s correlation coefficients (*r^2^*) are given in the plots.

**Figure S13:**
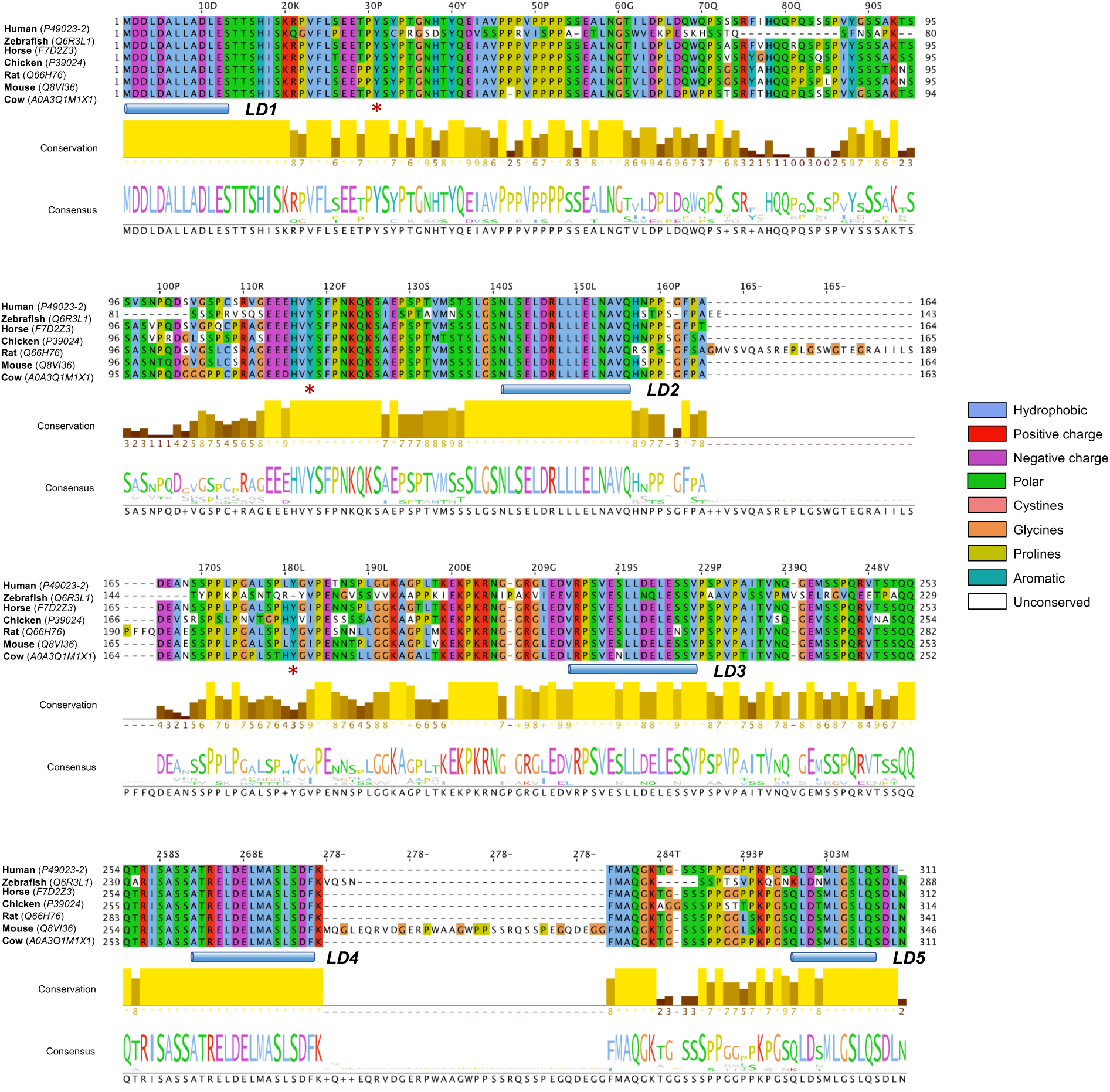
Sequence conservation of PXN from multiple species. Multiple sequence alignment of PXN from different species showing the conservation status of the intrinsically disordered regions. Sequences are labeled by the name of the species in bold, followed by the Uniprot ID in brackets. Residue positions are colored according to amino acid type as indicated in the figure. Conservation scores are represented by the yellow bar plot, where lighter shades of yellow indicate more conserved regions. The consensus sequence is highlighted using amino acid logo. The positions of the LD motifs are marked by the blue cylinders. Locations of the three tyrosine phospho-sites are marked by red star symbols.

**Figure S14:**
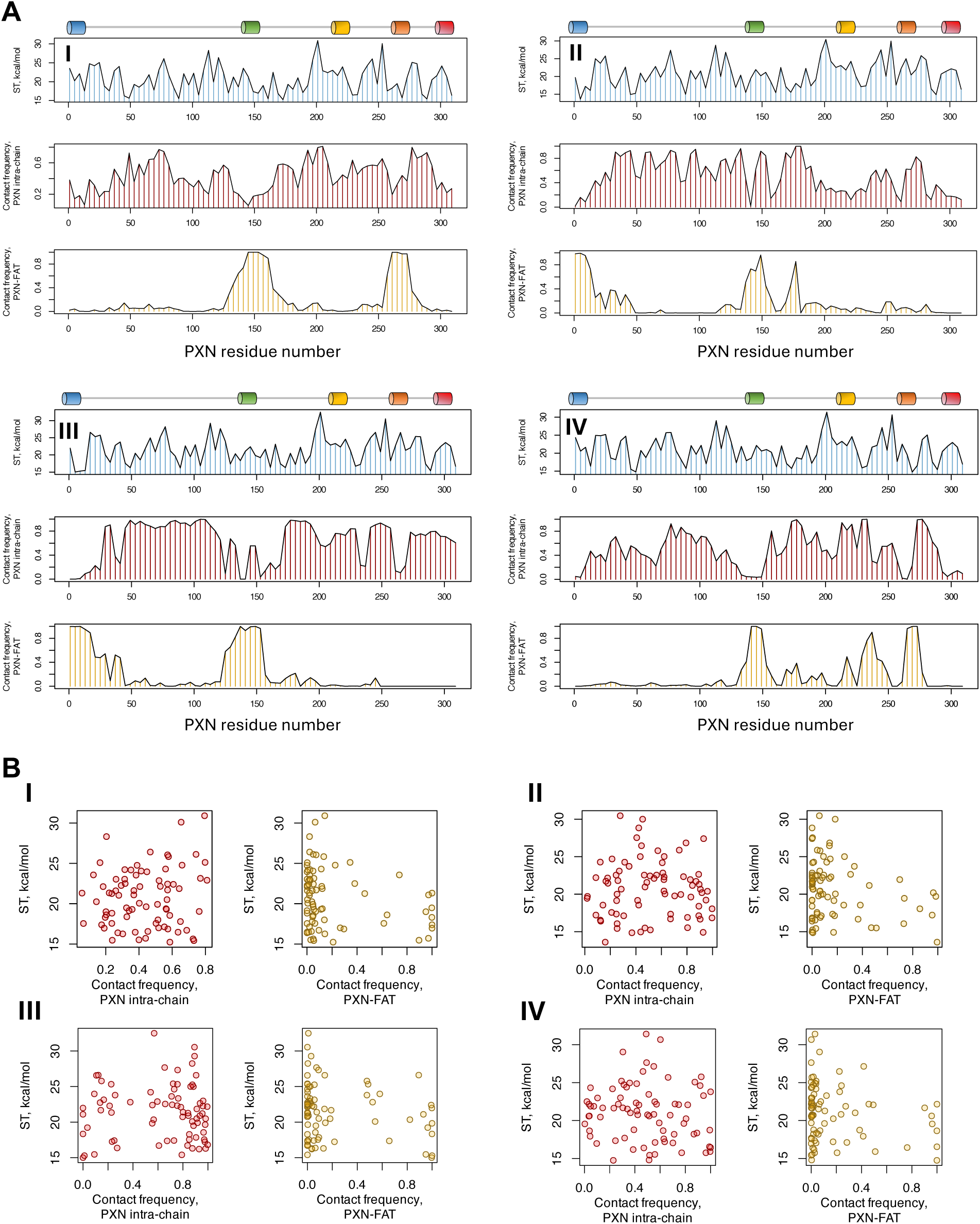
Conformational entropy of the disordered linker regions in PXN. (**A**) Residue-wise entropy (blue), intra-chain (red) and PXN-FAT (yellow) contact frequencies are plotted as function of PXN sequence for all four configurations. Entropy unit is the same as described in Fig. 10. Locations of the LD helices are shown above each plot using the same color convention that is throughout the manuscript. (**B**) Intra-chain (red dots) and PXN-FAT (yellow dots) contact frequencies for PXN residues plotted against their entropy values separately for the four configurations.

**Table S1:** Statistics for 10 best FAT structures.

|  |  |
| --- | --- |
| *A. Experimental chemical shift inputs* |  |
| <sup>13</sup>C<sub>α</sub> | 113 |
| <sup>13</sup>C<sub>β</sub> | 110 |
| <sup>13</sup>CO | 109 |
| <sup>15</sup>N | 95 |
| <sup>1</sup>H<sub>N</sub> | 95 |
| <sup>1</sup>H<sub>α</sub> | 84 |
| *B. RMSDs to the mean structure (Å)* |  |
| Over all residues <sup>a</sup> |  |
| Backbone atoms | 2.17 ± 0.52 |
| Heavy atoms | 2.72 ± 0.59 |
| Secondary structures <sup>b</sup> |  |
| Backbone | 1.86 ± 0.50 |
| Heavy atoms | 2.41 ± 0.56 |
| *C. Measures of structure quality* |  |
| Ramachandran distribution (%) <sup>c</sup> |  |
| Most favored | 96.2 ± 2.1 |
| Additionally allowed | 3.8 ± 2.1 |
| Generously allowed | 0.0 ± 0.0 |
| Disallowed | 0.0 ± 0.0 |
| *D. PDB/BMRB codes* |  |
| PDBDEV | 00000391 |
| BMRB | 51556 |

<sup>a</sup> Residues 918-1041.
<sup>b</sup> The secondary elements used were as follows: residues 920-940 (α1), 947-975 (α2), 977-1006(α3), 1012-1040 (α4).
<sup>c</sup> Ramachandran distributions were measured with Procheck.

